# Strain-specific virulence signatures of *Campylobacter jejuni* associated with watery versus bloody diarrhea in the neonatal piglet model

**DOI:** 10.1101/2025.04.25.650680

**Authors:** Jennifer M. Bosquez, Ben Pascoe, Craig T. Parker, Kerry K. Cooper

**Affiliations:** School of Animal and Comparative Biomedical Sciences, University of Arizona, Tucson, AZ 85721, USA; Ineos Oxford Institute for Antimicrobial Research, Department of Biology, University of Oxford, Oxford, United Kingdom; Faculty of Veterinary Medicine, Chiang Mai University, Chiang Mai, Thailand; Produce Safety and Microbiology Research Unit, Agricultural Research Service, United States Department of Agriculture, Albany, CA 94710, USA; BIO5 Institute, University of Arizona, Tucson, AZ 85719, USA

**Keywords:** watery diarrhea, neonatal piglet model, inflammatory diarrhea, bloody diarrhea, virulence, phenotypes

## Abstract

*Campylobacter jejuni* is the leading cause of bacterial gastroenteritis globally, presenting with either watery or bloody/inflammatory diarrhea. Despite this clinical variability, the underlying mechanisms driving these distinct diarrheal manifestations remain unclear. The neonatal piglet model provides a unique opportunity to investigate these differences using strains with defined disease manifestations. We evaluated ten *C. jejuni* strains previously shown to cause consistent diarrheal manifestations in neonatal piglets: five strains associated with watery diarrhea and five associated with bloody/inflammatory diarrhea. The study found significant strain to strain variation among most assays, but overall strains associated with bloody/inflammatory diarrhea exhibited significantly higher levels of epithelial cell invasion, intracellular survival, macrophage survivability, and disruption of intestinal barrier integrity compared to watery diarrhea strains. Notably, both groups had similar levels of adherence, produced similar levels of cytolethal distending toxin (CDT) and elicited comparable IL-8 responses. While both groups facilitated translocation of commensal *E. coli*, only bloody/inflammatory strains caused increased paracellular permeability, as shown by TEER and FITC-dextran assays. This study reveals that there is significant variation in the virulence phenotype of *C. jejuni* strains and identifies several characteristics consistently associated with a specific diarrheal manifestation group. These findings provide new insights into the strain-specific pathogenic mechanisms of *C. jejuni*, with implications for understanding host-pathogen interactions and informing targeted diagnostic and therapeutic strategies.

**Author Summary:** *Campylobacter jejuni* is one of the most common bacterial causes of diarrhea worldwide, affecting millions each year. Infected individuals may experience either watery or bloody/inflammatory diarrhea, but the reasons why different strains cause different types of illness have remained a mystery. In this study, we used a neonatal piglet model, which accurately mimics human diarrheal disease, to investigate why certain *C. jejuni* strains cause more severe symptoms than others. We compared strains known to cause either watery or bloody diarrhea and found key differences in their ability to invade intestinal cells, survive inside immune cells, and damage the intestinal barrier. While both groups of strains could trigger inflammation and translocate *E. coli* through the intestinal lining, only the strains linked to bloody diarrhea caused significant barrier disruption and had increased intracellular survival. These findings reveal that different *C. jejuni* strains follow distinct paths during infection, which helps explain why they produce different symptoms. Unraveling these differences improves our overall understanding of *Campylobacter* pathogenesis that could lead to improved diagnostics and/or treatments to reduce the burden of campylobacteriosis around the world.

## Introduction

*Campylobacter jejuni* is the leading cause of bacterial gastroenteritis worldwide, responsible for over 550 million cases annually[1]. In the United States alone, it affects more than 1.9 million people per year, imposing significant economic and public health burdens[2]. While typically self-limiting, *C. jejuni* infection can lead to long-term sequelae such as post-infectious irritable bowel syndrome (PI-IBS), reactive arthritis, and Guillain-Barré syndrome[3–5]. Increasing evidence also links *Campylobacter* to growth stunting in children from low- and middle-income countries due to chronic enteric infections[6–8].

Clinically, *C. jejuni* infection presents as watery or bloody/inflammatory diarrhea, with up to 27% of U.S. cases involving visible blood in stools[9]. Bloody/inflammatory diarrhea suggests deeper intestinal invasion and inflammation, often associated with more severe outcomes[10]. Despite decades of reporting this clinical distinction, the mechanisms driving these divergent diarrheal manifestations remain poorly understood. Several virulence factors have been proposed to influence disease severity, including enterotoxin production, cytotoxins like cytolethal distending toxin (CDT), and bacterial invasiveness[11–13]. However, no studies have definitively linked these traits to specific diarrheal outcomes. Clinical data are often confounded by subjective patient reporting and variable strain backgrounds, complicating direct comparisons. Additionally, many *Campylobacter* studies rely heavily on just a few clinical *C. jejuni* strains, most commonly NCTC11168, 81-176, or 81116[14, 15], which may unintentionally bias findings, given the well-documented heterogeneity in virulence factors across *C. jejuni* strains virulence factors[16–18].

The neonatal piglet model provides a robust platform for studying *C. jejuni*-induced diarrhea[19]. Unlike murine models, piglets exhibit human-like disease and can consistently replicate both watery and bloody/inflammatory diarrheal manifestations following infection with specific *C. jejuni* strains[20, 21]. Additionally, piglets are recognized as strong alternatives to both canines or non-human primates for non-rodent evaluations that are required by regulatory agencies[22, 23], and also have intestinal functions physiologically similar to humans[24–26]. Utilization of this model allows for controlled, strain-specific comparisons with minimized host variation. Healthy piglets typically do not develop disease when exposed to *C. jejuni*[27]. However, colostrum-deprived neonatal piglets consistently develop clinical symptoms such as large intestinal lesions, inflammation, and diarrhea lasting up to six days depending on the *C. jejuni* strain used for infection[28]. Similarly, gnotobiotic piglets infected with *C. jejuni* also display symptoms of diarrhea and sloughing of epithelial cells from the mucosa in the cecum and colon closely resembling human campylobacteriosis pathologies[20, 29]. Despite these advantages, the use of piglets in *C. jejuni* research remains limited due to the high costs and specialized housing requirements, leading many researchers to favor more accessible models such as mice, ferrets, guinea pigs, or rats.

In this study, we leveraged the neonatal piglet model to investigate phenotypic differences between *C. jejuni* strains that cause watery diarrhea compared to those that cause bloody/inflammatory diarrhea. Using ten strains classified by the resultant diarrheal manifestation, we assessed virulence traits including invasion, intracellular survival, epithelial barrier disruption, and immunostimulation. Our goal was to evaluate these virulence and pathogenicity traits that may relate to the differences in diarrheal presentation, advancing our understanding of *C. jejuni* pathogenesis and informing new approaches to diagnosis or intervention.

## Materials and Methods

### Bacterial Culturing and Growth

*C. jejuni* strains used in this study are listed in **Table 1**. All strains were grown and maintained on Mueller-Hinton agar plates supplemented with 5% defibrinated bovine blood (MHB; Becton, Dickinson and Company Difco) for 48 hours at 37°C with 10% CO_2_. Strains for assays that required liquid culture were grown in Mueller-Hinton broth (MH; Becton, Dickinson and Company Difco) with constant shaking (100 rpm) for 16-18 hours at 37°C with 10% CO_2_. Assay control strains *Escherichia coli* DH5α and *Salmonella enterica* subspecies *enterica* serovar Typhimurium str. RM6835 were both grown aerobically either on Luria-Bertani (LB) agar (Fisher Bioreagents) at 37°C for 24 hours or in LB broth (Fisher Bioreagents) with constant shaking (100 rpm) at 37°C for 24 hours. For selective isolation of *Campylobacter* in select assays, *C. jejuni* strains were cultured on *Campylobacter* CEFEX agar, a modified MH agar with the addition of vancomycin (20 mg/L), polymyxin B (0.35 mg/L), trimethoprim (26.4 mg/L), sodium cefoperazone (20 mg/L) and amphotericin B (10 mg/L).

**Table 1.** Campylobacter jejuni strains utilized in this study.

| Strain | Source | Sequence Type (ST) | Clonal Complex | Diarrheal Manifestation |
| --- | --- | --- | --- | --- |
| NCTC11168 | Clinical | ST43 | ST-21 | Bloody and/or Inflammatory |
| 81-176 | Clinical | ST604 | ST-42 | Bloody and/or Inflammatory |
| M129 | Clinical | ST353 | ST-353 | Bloody and/or Inflammatory |
| A9a | Poultry | ST2827 | ST-460 | Bloody and/or Inflammatory |
| D33a | Poultry | ST459 | ST-42 | Bloody and/or Inflammatory |
| RM1221 | Poultry | ST354 | ST-354 | Watery |
| S3 | Poultry | ST354 | ST-354 | Watery |
| D42a | Poultry | ST21 | ST-21 | Watery |
| A14a | Poultry | ST1212 | ST-607 | Watery |
| A13a | Poultry | ST1212 | ST-607 | Watery |

### Cytolethal Distending Toxin (CDT) Assay

CDT activity was measured using a modified protocol from Abuoun et al[30]. Briefly, *C. jejuni* strains were grown overnight on MHB plates at either 37°C or 42°C under microaerobic conditions (10% CO₂). Bacterial growth was harvested by resuspending the entire plate in 5 ml of *Brucella* broth (Becton, Dickinson and Company, BBL). The suspension was centrifuged at 10,000 × g for 20 minutes at 4°C. The supernatant was discarded, and the bacterial pellet was resuspended in 3 ml of Eagle’s Minimum Essential Medium (EMEM; Corning) supplemented with 10% fetal bovine serum (FBS; Corning). The OD₆₀₀ of the suspension was adjusted to 1.0 using EMEM +10% FBS. Lysates were prepared by sonicating the suspension on ice using a QSonica sonicator (20-second pulses, 20-second rests, five cycles total). The lysate was centrifuged at 10,000 × g for 20 minutes at 4°C, and the supernatant was filter-sterilized using a 0.22 µm syringe filter (Fisher Scientific) to remove viable bacteria. Sterile lysates were serially diluted two-fold (1:1 to 1:128) in EMEM + 10% FBS. HeLa cells (ATCC CCL-2) were seeded into 96-well plates at 2 × 10⁴ cells/ml in 100 µl of EMEM + 10% FBS and incubated for 3 hours at 37°C with 5% CO₂. Subsequently, 100 µl of each dilution of filtered cell lysate was added to the HeLa cells. Plates were incubated at 37°C with 5% CO₂ for five days. Following incubation, cells were fixed with 10% formalin and stained with 0.8% crystal violet. The CDT titer was defined as the reciprocal of the highest dilution showing >50% cell distension. *E. coli* DH5α was used as a negative control. Each strain was tested in three biologically independent assays (n = 3).

### Attachment Assays

Attachment was assessed using three epithelial cell lines: non-polarized INT407 (ATCC CCL-6), and polarized Caco-2 BBE (ATCC HTB-37) and T84 (ATCC CCL-248) cells. Cell lines were seeded in 24-well tissue culture-treated plates (1 ml per well at 1 × 10⁶ cells/ml) and incubated for 18 hours at 37°C with 5% CO₂. *Campylobacter* cultures were grown overnight in MH broth with shaking (100 rpm) at 37°C in 10% CO₂. The OD₆₀₀ was adjusted to 0.1 in the appropriate medium for each cell line: EMEM + 10% FBS for INT407, EMEM + 20% FBS for Caco-2 BBE, and DMEM/F-12 + 5% FBS for T84. Each well was washed twice with sterile PBS, then inoculated with 1 ml of *C. jejuni* inoculum (approximately 1 × 10⁸ CFU, multiplicity of infection (MOI) 100). To enhance bacterial contact with host cells, plates were centrifuged at 600 × g for 5 minutes then incubated for 1 hour at 37°C with 5% CO₂. Wells were then washed twice with PBS to remove non-adherent bacteria. Host cells were lysed by adding 1 ml of 0.1% Triton X-100 in PBS and incubating for 10 minutes at room temperature. Lysates were serially diluted and plated on MHB agar for CFU enumeration after incubation at 37°C in 10% CO₂ for 48 hours. Attachment percentage was calculated as: (CFU recovered / CFU inoculated) × 100. Each strain was tested in technical duplicates in three independent experiments (n = 3). Negative controls included two wells inoculated with only sterile PBS. *E. coli* DH5α was used as a positive control. Due to insufficient T84 cell growth, *C. jejuni* strain 81-176 was unable to be included in attachment assays using this line.

### Invasion assays

Enterocyte invasion was assessed using INT407, Caco-2 BBE, and T84 cells, following the same seeding and infection conditions described for attachment assays. After overnight growth of *C. jejuni* in MH broth at 37°C with 10% CO₂ and shaking (100 rpm), cultures were diluted to OD₆₀₀ = 0.1 in the appropriate medium for each cell line to achieve a MOI of 100. Each well of the 24-well plates was washed twice with PBS and then inoculated with 1 ml of the bacterial suspension. Plates were centrifuged at 600 × g for 5 minutes and incubated for 3 hours at 37°C with 5% CO₂ to allow for bacterial invasion. Following incubation, wells were washed twice with PBS, and 1 ml of cell line-specific medium containing gentamicin (250 µg/ml; Acros Organics) was added to each well. Plates were incubated for an additional hour to eliminate extracellular bacteria. After gentamicin treatment, wells were washed twice with PBS and then lysed with 1 ml of 0.1% Triton X-100 in PBS. The lysates were serially diluted and plated on MHB agar, which was incubated at 37°C with 10% CO₂ for 48 hours. The percent invasion was calculated as: (CFU recovered after gentamicin / CFU in initial inoculum) × 100. Each strain was assessed in technical duplicates in three biologically independent experiments (n = 3). *Salmonella* Typhimurium strain RM6835 was used as a positive control, and *E. coli* DH5α as a negative control. Every assay included two uninoculated wells and were processed exactly as the positive and negative controls. Due to insufficient T84 cell growth, *C. jejuni* strain 81-176 was unable to be included in invasion assays using this cell line.

### Macrophage Survivability

The ability of the different *C. jejuni* strains to survive intracellularly in macrophages was assessed using J774A.1 mouse macrophages (ATCC TIB-67). Cells were seeded into 24-well plates at 1 × 10⁶ cells/ml in 1 ml per well and incubated for 18 hours at 37°C with 5% CO₂ to establish monolayers. Overnight MH broth cultures of each *C. jejuni* strain were prepared, adjusted to OD₆₀₀ = 0.1 (∼1 × 10⁸ CFU/ml) in DMEM + 10% FBS (J7774A.1 culture media), and added to each well at an MOI of 100. Two wells were left uninoculated as negative controls. Plates were incubated at 37°C with 5% CO₂ for 24, 48, or 72 hours. At each time point, wells were washed twice with PBS and then treated with 1 ml of gentamicin (250 µg/ml in DMEM + 10% FBS) for 1 hour to eliminate extracellular bacteria. After treatment, wells were washed again, and macrophages were lysed by adding 1 ml of 0.1% Triton X-100 in PBS. Lysates were serially diluted and plated on MHB agar, incubated at 37°C with 10% CO₂ for 48 hours, and colonies enumerated. Each strain was tested in two technical replicates across three independent biological assays (n=3). *C. jejuni* strain 81-176 was included as a positive control due to its known interaction with macrophages[31].

### IL-8 ELISA and 24-hour Intracellular Survivability

T84 cells were seeded into 24-well plates at 5.5 × 10⁵ cells/well and cultured for approximately 7 days until reaching 100% confluence. *C. jejuni* strains were grown overnight in MH broth and diluted to OD₆₀₀ = 0.1 in DMEM/F-12 with 5% FBS (T84 medium). Each well was infected with 1 ml of bacterial suspension (∼1 × 10⁸ CFU, MOI 100) and incubated at 37°C with 5% CO₂ for 3 hours. Following infection, wells were washed with PBS and treated with 1 ml of gentamicin (250 µg/ml in T84 medium) for 1 hour. Wells were then washed again, fresh medium was added, and plates were incubated for 24 hours at 37°C with 5% CO₂. After 24 hours, media from each well was collected, centrifuged at 12,000 × g for 1 minute, and the supernatant stored at –80°C for cytokine analysis. Human IL-8 levels were measured using the BD OptEIA™ ELISA kit (BD Biosciences) per the manufacturer’s instructions. To assess intracellular bacterial survival, epithelial cells were lysed after supernatant removal by adding 1 ml of 0.1% Triton X-100 in PBS. Lysates were serially diluted and plated on MHB agar. Plates were incubated at 37°C in 10% CO₂ for 48 hours, and colonies were counted. Each strain was evaluated in two technical replicates per assay, with three independent biological replicates (n = 3). Due to insufficient T84 cell growth, *C. jejuni* strain 81-176 was unable to be included in IL-8 and intracellular assays using this cell line.

### Transepithelial Electrical Resistance (TEER)

To evaluate epithelial barrier integrity, transepithelial electrical resistance (TEER) was measured using Caco-2 BBE cells. Cells were seeded at 2 × 10⁵ cells per well into 0.4 μm pore-size Transwell inserts (Falcon, Corning) placed in 24-well tissue culture-treated plates. Cells were cultured in EMEM with 20% FBS at 37°C with 5% CO₂ until a confluent monolayer formed (TEER > 250 Ω·cm²), typically by day 14. Monolayer integrity was assessed microscopically and with an EVOM2 volt/ohm meter (World Precision Instruments). Following confluence, cells were maintained for an additional 120 hours to promote tight junction maturation. Prior to infection, TEER was recorded, and a cell count was performed using a hemocytometer to calculate the bacterial concentration needed for MOI 100. *C. jejuni* cultures were prepared by growing strains overnight in MH broth, adjusting to OD₆₀₀ = 0.1, and diluting in Caco-2 BBE medium. From each Transwell, 50 µl of media was removed from the apical compartment and replaced with 50 µl of the corresponding *C. jejuni* inoculum (∼5 × 10⁶ CFU; MOI 100). Each strain was tested in duplicate Transwells, with two uninoculated wells serving as negative controls (sterile medium only). Infected plates were incubated at 37°C with 5% CO₂ for 72 hours. TEER measurements were taken at 24, 48, and 72 hours using Millicell ERS-2 electrodes (Millipore). Electrodes were disinfected and rinsed between each well using the following sequence: 10% bleach, sterile water (×2), 70% ethanol, sterile water (×2), and pre-warmed media. Each *C. jejuni* strain was tested in duplicates during three independent biological experiments (n = 3).

### FITC-Dextran Translocation

To assess paracellular permeability to small molecules, a 4-kDa fluorescein isothiocyanate (FITC)-dextran probe (Millipore Sigma) was applied to mature Caco-2 BBE monolayers grown on 0.4 µm Transwell inserts in phenol red-free MEM (Gibco). Cells were seeded at 2 × 10⁵ cells per insert and cultured to confluence (TEER > 250 Ω·cm²), then maintained for an additional 120 hours to allow tight junction maturation. Overnight *C. jejuni* cultures were adjusted to OD₆₀₀ = 0.1 (∼5 × 10⁶ CFU) and used to inoculate the apical compartments (50 µl per insert, MOI 100). Two wells were left as negative controls (sterile medium only). After 72 hours of incubation at 37°C with 5% CO₂, 100 µl of media was removed from the apical compartment and replaced with 100 µl of 4-kDa FITC-dextran solution (10 mg/ml in PBS). Plates were incubated for 3 hours, then 100 µl was collected from the basolateral compartment and transferred to a black walled 96-well plate for fluorescence measurement. Fluorescence was measured using a SpectraMax M2 plate reader (Molecular Devices) with excitation at 490 nm and emission at 530 nm. Relative fluorescence units (RFUs) were blanked against sterile Caco-2 BBE medium. Each *C. jejuni* strain was tested in three biological experiments with two technical replicates each (n = 3).

### Commensal Bacteria Translocation

The ability of *C. jejuni* to disrupt the epithelial barrier and promote translocation of non-invasive commensal bacteria was assessed using *E. coli* DH5α as a model commensal. Two experimental approaches were performed: (1) a 6-hour co-inoculation and (2) a 72-hour pre-infection with *C. jejuni* followed by 3-hour *E. coli* exposure. For both approaches, Caco-2 BBE cells were seeded at 1.5 × 10⁵ cells/ml into 3 µm pore-size Transwell inserts and cultured to confluence (TEER > 250 Ω·cm²), with daily monitoring of TEER.

#### 6-hour co-culture

Overnight cultures of *C. jejuni* and *E. coli* DH5α were grown and adjusted to OD₆₀₀ = 0.1 (∼5 × 10⁶ CFU). The apical compartment was inoculated with 50 µl of each strain (MOI 100 per organism). After 6 hours of incubation, media from the basal compartment was collected and plated on MacConkey agar for *E. coli* and CEFEX agar for *C. jejuni*. CEFEX agar was incubated at 37°C with 10% CO₂ for 48 hours; MacConkey plates were incubated aerobically at 37°C for 24 hours. Each strain was tested in three independent biological experiments with two technical replicates each (n = 3).

#### 72-hour pre-infection + 3-hour E. coli exposure

Caco-2 BBE cells were infected with *C. jejuni* as described for the TEER assay. After 72 hours, 50 µl of *E. coli* DH5α (MOI 100) was added to the apical compartment and incubated for an additional 3 hours. Basolateral media was collected and plated as described above. Each strain was tested in three independent biological experiments with two technical replicates per experiment (n = 3). To assess intracellular *C. jejuni* and *E. coli*, the apical compartment was treated with gentamicin (250 µg/ml) for 1 hour, washed, and lysed with 0.1% Triton X-100. Lysates were plated on CEFEX and MacConkey agar for quantification. Translocation efficiency was calculated as: (CFU in basal compartment / CFU inoculated) × 100.

### Statistical Analysis

Statistical analyses were performed using GraphPad Prism (v10.4.1) for comparisons between diarrheal manifestations or RStudio (v2022.07.1) for comparisons between strains. All data were tested for normality (Shapiro–Wilk test) and equal variance (Levene’s test). For comparisons between strains associated with watery and bloody/inflammatory diarrhea (n = 15 per group), the non-parametric Mann–Whitney U test was applied. For comparisons across multiple strains or conditions, non-parametric Kruskal–Wallis tests with Dunn’s post hoc test were used. Compact Letter Display (CLD) notation was used to visualize the results of the pairwise comparisons. Strains that do not share a letter are significantly different from each other (p < 0.05), while strains sharing the same letter are not significantly different. Letters were assigned using the multcompView package in R, based on adjusted p-values. Statistical significance was defined as p ≤ 0.05. Each assay included a minimum of three independent biological replicates per strain (n = 3 per group), with technical replicates as noted. Data are presented as mean ± SEM unless otherwise stated.

## Results

### C. jejuni strains associated with bloody/inflammatory diarrhea exhibit increased invasion and intracellular survival in epithelial cells

To assess whether *C. jejuni* strains associated with distinct diarrheal phenotypes exhibit differences in virulence potential both on a strain-to-strain level and between groups, we compared five strains associated with watery diarrhea and five associated with bloody/inflammatory diarrhea (**Table 1**). All ten strains have previously been investigated in the neonatal piglet model to confirm their diarrheal manifestations *in vivo* (**Fig 1A-B**). Invasion assays were conducted using INT407, Caco-2 BBE and T84 cells. Significant differences were observed between several strains, notably, A9a was a significantly better invader of Caco-2 BBE and T84 cells than most other strains, particularly the watery diarrhea associated strains (Note: 81-176 was not tested with T84 cells), whereas all five of the watery diarrhea associated strains were poor invaders regardless of the cell line tested (**Fig 2A, Supplementary Fig S1A, Supplementary Fig S2A**). Interestingly, no significant differences were observed on a strain to strain level using INT407 cells. Invasion assays conducted in polarized Caco-2 BBE cell revealed significantly greater invasion by bloody/inflammatory associated strains compared to watery diarrhea strains (p = 0.0014; **Fig 2B**) Similar trends were observed using non-polarized INT407 cells (p = 0.0006; **Supplementary Fig S1B**) and T84 cells (p = 0.0063; **Supplementary Fig S2B**).

**Fig 1.**
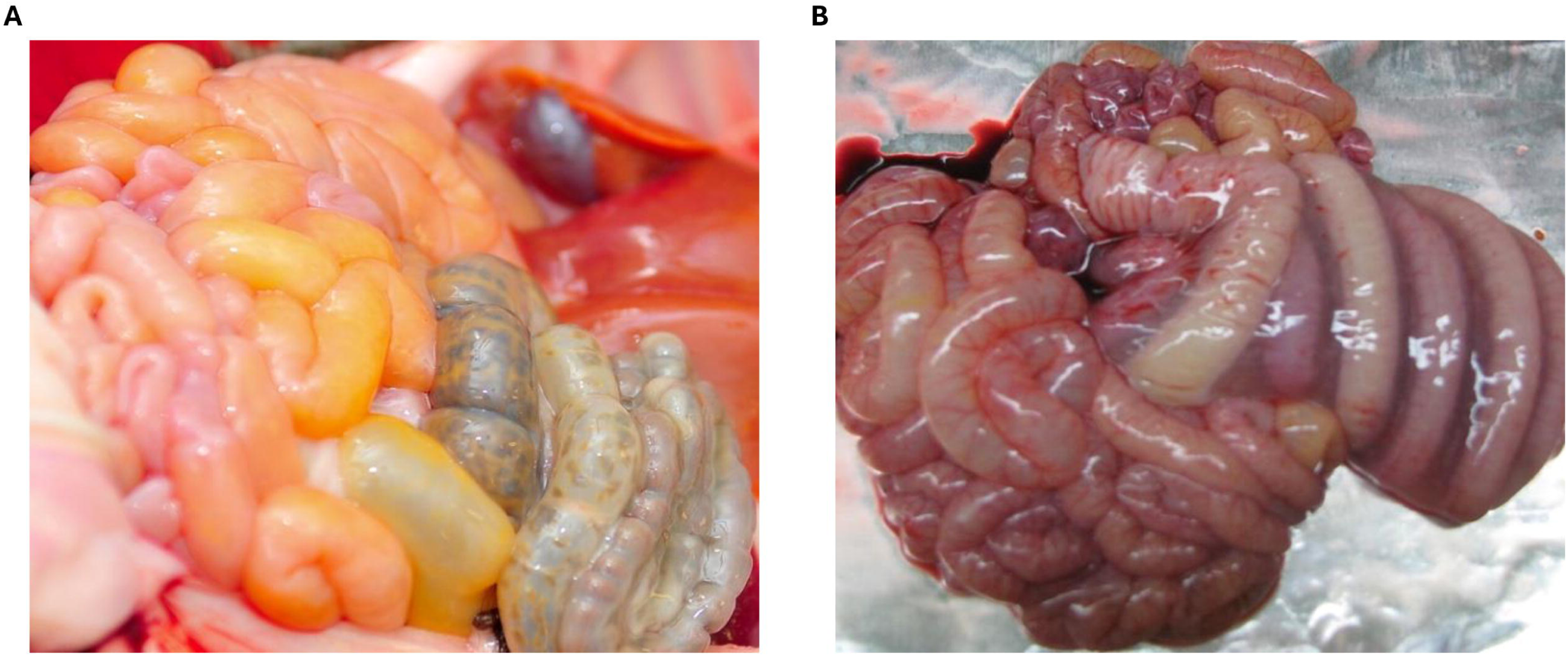
Gross pathology associated with different diarrheal outcome caused by different strains of *Campylobacter jejuni* based on previous challenge in colostrum deprived neonatal piglet model. (A) Gross pathology of the intestinal tract of neonatal piglets infected with *C. jejuni* strains associated with watery diarrhea manifestation. (B) Gross pathology of the intestinal tract of neonatal piglets infected with *C. jejuni* strains associated with bloody and/or inflammatory diarrhea manifestation. Piglet studies were conducted prior to this study and were described previously in Law et al and Cooper et al[21, 35].

**Fig 2.**
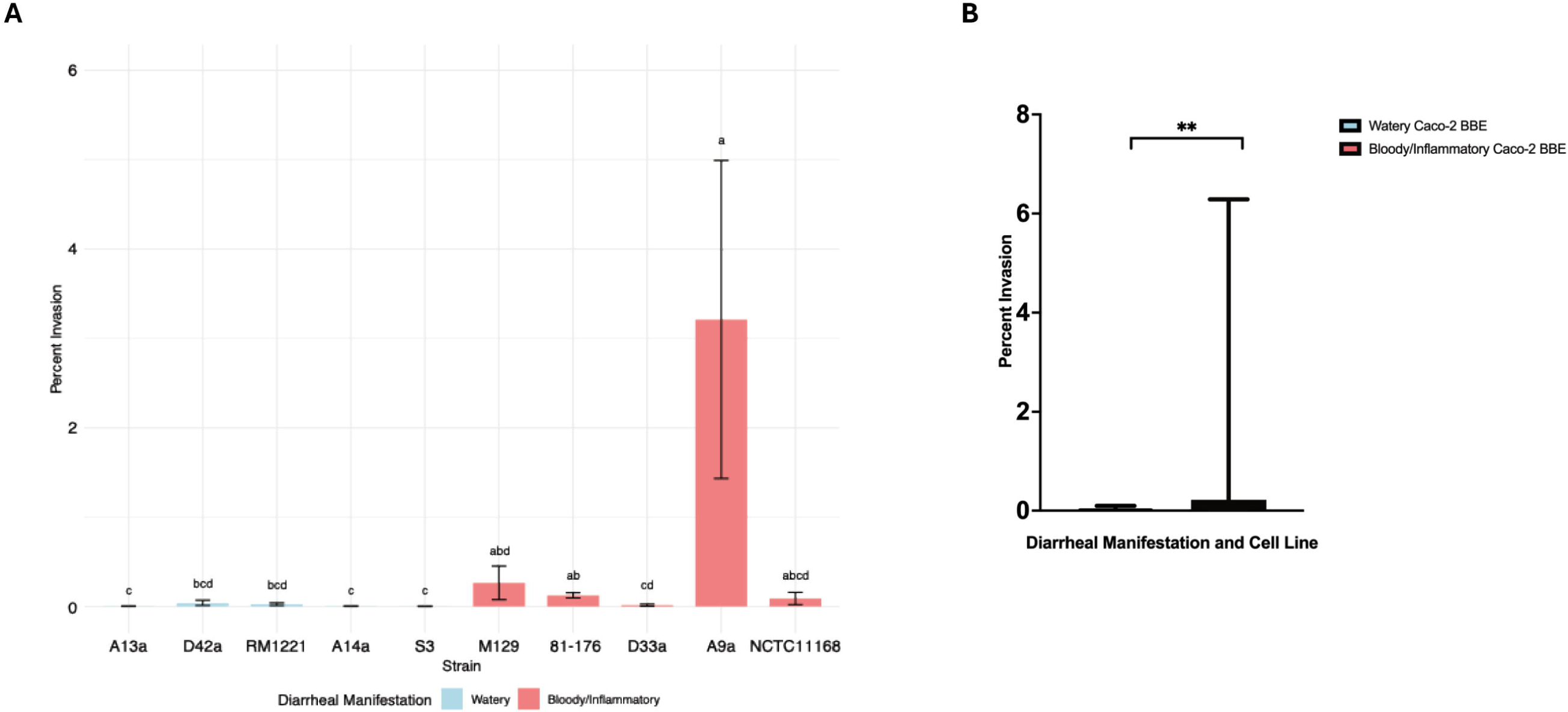
Average percent invasion of intestinal epithelial cell line Caco-2 BBE. (A) Invasion across all ten *C. jejuni* strains. Bars labeled with different letters indicate statistically significant differences (p < 0.05) based on Dunn’s post hoc test following a Kruskal-Wallis analysis. Shared letters denote groups that are not significantly different from each other. (B) Invasiveness of Caco-2 BBE cells by *C. jejuni* strains grouped by diarrheal manifestation (**, p = 0.0014).

To evaluate whether increased invasion correlated with improved intracellular persistence, we measured viable intracellular bacteria at 24 hours post-infection in T84 cells. Strain to strain variation was observed among bloody/inflammatory strains M129 and A9a, where they displayed significantly higher intracellular survival than most other strains (**Fig 3A**). As a group, bloody/inflammatory diarrhea associated strains exhibited significantly higher intracellular survival than watery diarrhea associated strains (p = 0.0007; **Fig 3B**).

**Fig 3.**
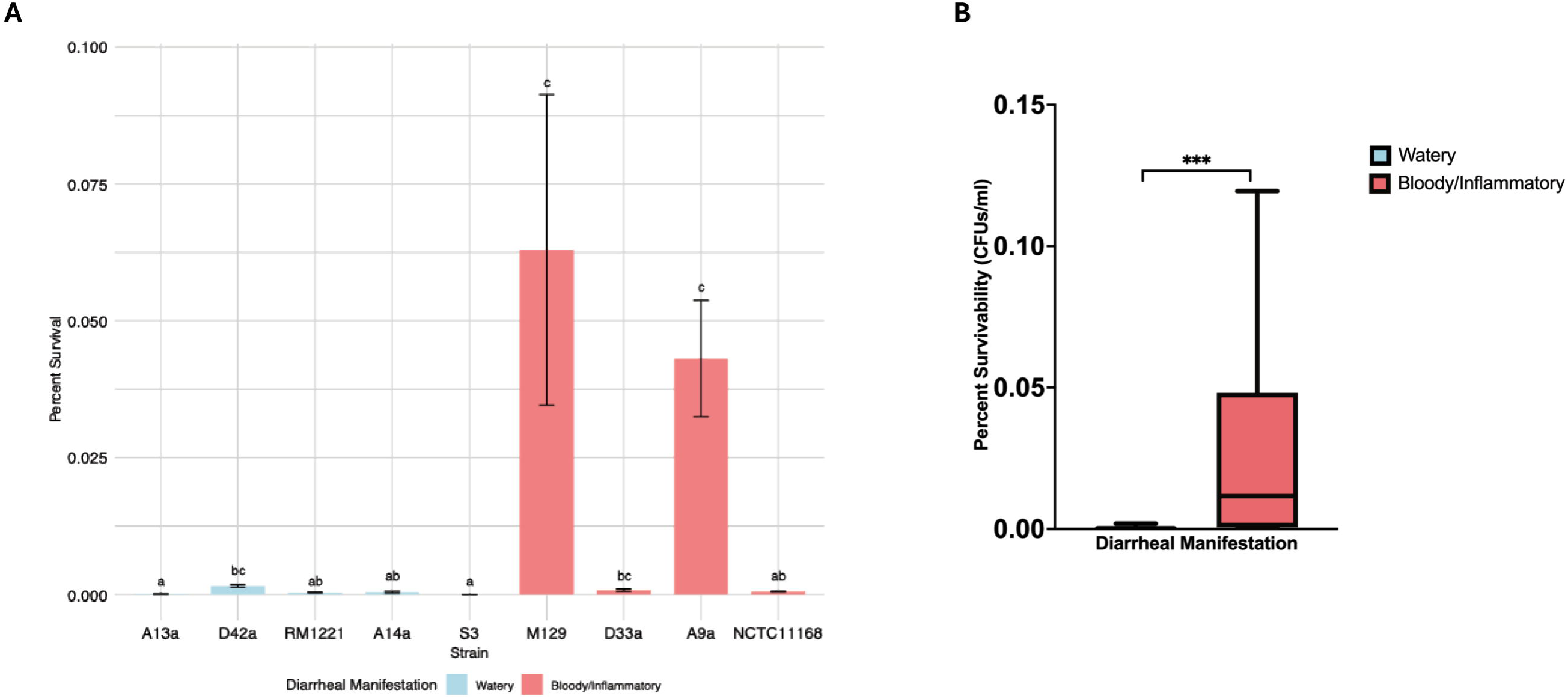
Intracellular survival over 24 hours in T84 cells. (A) Average percent survival within T84 cells by strain. Bars labeled with different letters indicate statistically significant differences (p < 0.05) based on Dunn’s post hoc test following a Kruskal-Wallis analysis (n = 3). Shared letters denote groups that are not significantly different from each other. (B) Percent T84 24-hour intracellular survival by *C. jejuni* strains grouped by diarrheal manifestation (***, p = 0.0007). Strain 81-176 is not included in this data set.

### Similar attachment of C. jejuni strains across intestinal cell lines

To determine whether differences in invasion were due to upstream differences in adherence, we assessed attachment efficiency after a 1-hour infection of epithelial cell lines INT407, Caco-2 BBE and T84. Few significant differences were observed between strains across INT407 (**Fig 4A**), Caco-2 BBE cells (**Fig 4C**) and T84 cells (**Supplementary Fig S3A**). There were no significant differences in attachment between watery and bloody/inflammatory diarrhea associated strains, regardless of the cell line used: INT407 cells (p = 0.20; **Fig 4B**), Caco-2 BBE (p = 0.14; **Fig 4D**), T84 cells (p = 0.90; **Supplementary Fig S3B**).

**Fig 4.**
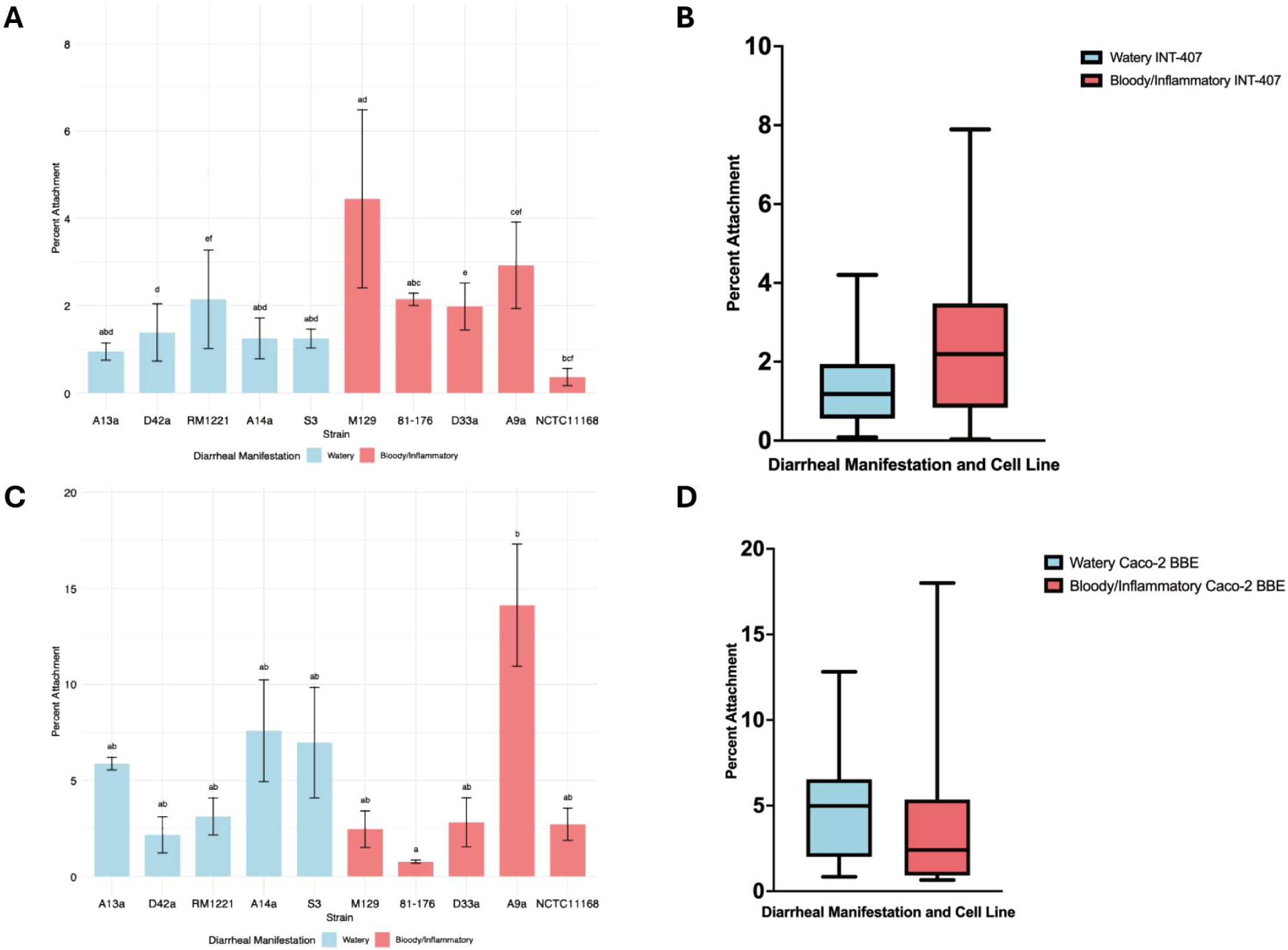
Adherence to intestinal epithelial cell lines. (A) Adherence of each *C. jejuni* strain to INT407 cells. Bars labeled with different letters indicate statistically significant differences (p < 0.05) based on Dunn’s post hoc test following a Kruskal-Wallis analysis (n = 3). Shared letters denote groups that are not significantly different from each other. (B) Adherence to INT407 cell by *C. jejuni* strains grouped by diarrheal manifestation (p = 0.20). (C) Adherence of each *C. jejuni* strain to Caco-2 BBE cells (n = 3). (D) Adherence to Caco-2 BBE cells by *C. jejuni* strains grouped by diarrheal manifestation (p = 0.14).

### CDT production does not distinguish between watery and bloody/inflammatory strains

All *C. jejuni* strains in this study produced detectable CDT activity, as measured by the HeLa cell distension assay. We did not observe any strain variation in CDT production between watery and bloody/inflammatory diarrhea associated strains at either 37°C or 42°C (**Supplementary Fig S4A)**. When compared as groups at 37°C, mean CDT titers were comparable between watery (geometric mean = 0.094) and bloody/inflammatory diarrhea associated strains (geometric mean = 0.086; p = 0.74; **Supplementary Fig S4B**). Due to the commensal relationship between poultry and *C. jejuni*, CDT production was also measured at the poultry internal body temperature of 42°C and here mean CDT titers were different between watery (geometric mean = 0.10) and bloody/inflammatory strains but not statistically significant (geometric mean = 0.041; p = 0.086; **Supplementary Fig S4B**).

### Bloody/inflammatory strains exhibit greater 24-hour intracellular survival in T84 cells but induce similar IL-8 responses

To evaluate whether differences in intracellular persistence affected epithelial inflammatory signaling, we quantified human IL-8 production in T84 cell supernatants at 24 hours post-infection and measured viable intracellular *C. jejuni* following gentamicin treatment and lysis. IL-8 production was not significantly different between strains but did show variability (**Supplementary Fig S5A**). Strain A9a induced the highest mean IL-8 levels (1202.04 pg/ml), whereas D33a produced the lowest (437.03 pg/ml), highlighting strain-level variability. Strains associated with bloody/inflammatory diarrhea demonstrated significantly greater intracellular survival at 24 hours than watery diarrhea strains (p = 0.0007; **Fig 3B**). Despite this, IL-8 secretion was not significantly different between groups: mean concentrations were 809.10 pg/ml for watery strains and 702.60 pg/ml for bloody/inflammatory strains (p = 0.61; Supplementary Fig S5B).

### Bloody/inflammatory strains more profoundly disrupt epithelial barrier function over time

To assess whether *C. jejuni* strains differ in their capacity to disrupt epithelial barrier integrity, we measured TEER in polarized Caco-2 BBE monolayers at 24-, 48-, and 72-hours post-infection. Across 24-, 48- and 72-hours, strains A14a, A13a and S3 had the lowest changes in TEER while strains A9a, M129 and 81-176 had the highest changes in TEER (**Fig 5A**). When grouped by diarrheal manifestation, bloody/inflammatory diarrhea associated strains caused a significantly greater reduction in TEER at all time points. At 24 hours, TEER decreased by an average 15.9% compared to 4.03% in watery diarrhea associated strains (p < 0.0001; **Fig 5B**). Differences remained significant at 48 hours (27.1% verses 10.7%; p < 0.0001) and 72 hours (43.4% verses 18.3%; p < 0.0001).

**Fig 5.**
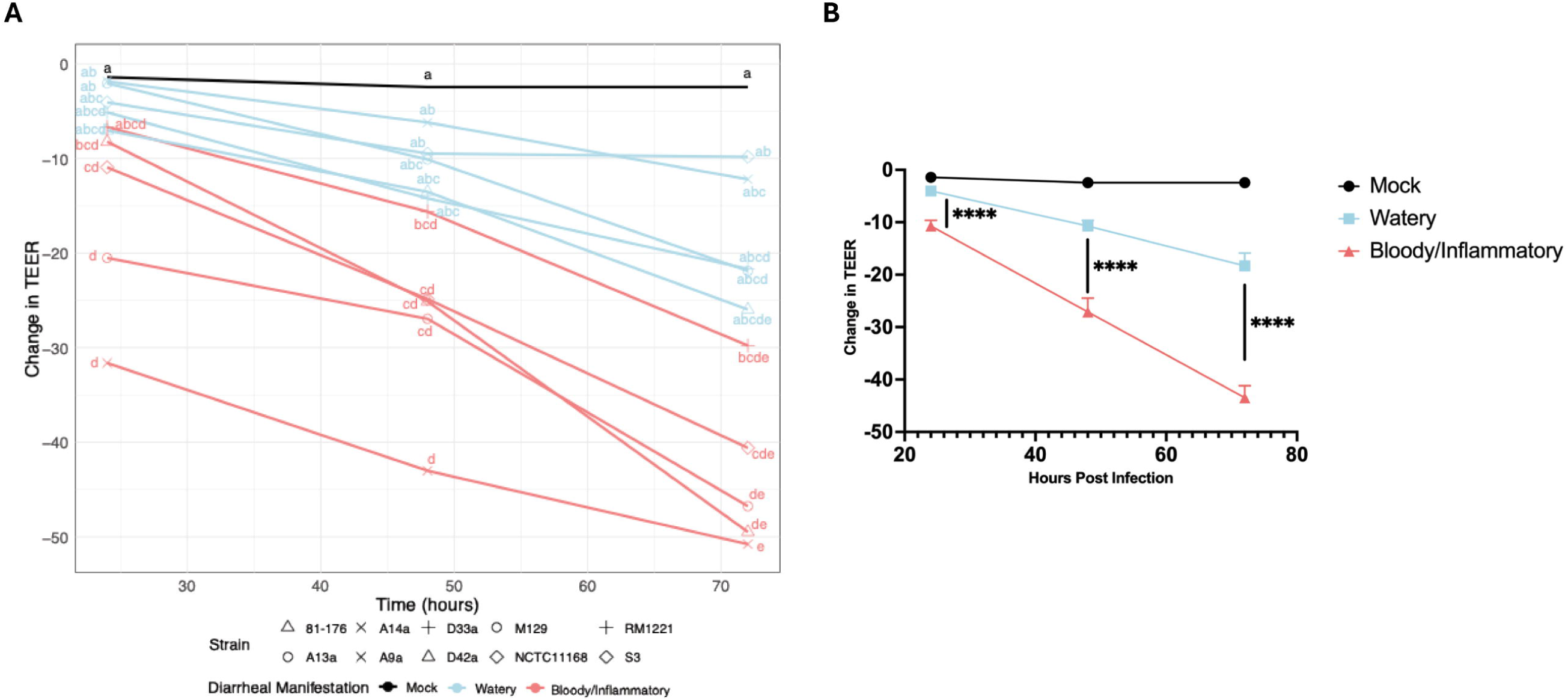
**Change in the transepithelial electrical resistance (TEER) over 24, 48 and 72 hours**. (A) Change in TEER for each *C. jejuni* strain over 24-, 48- and 72-hours. Those labeled with different letters indicate statistically significant differences (p < 0.05) based on Dunn’s post hoc test following a Kruskal-Wallis analysis (n = 3). Shared letters denote groups that are not significantly different from each other. (B) Change in TEER across *C. jejuni* strains grouped by diarrheal manifestation at 24 hours (****, p < 0.0001), 48 hours (****, p < 0.0001), and at 72 hours (****, p < 0.0001). Mock infection represents Caco-2 BBE cells that contained only sterile cell culture media.

### Bloody/inflammatory strains increase paracellular permeability to small molecules

To directly measure functional barrier permeability, we quantified apical-to-basolateral flux of 4-kDa FITC-dextran across polarized Caco-2 BBE monolayers following 72-hour infection. Strain-specific permeability values were observed to be highest among strain A9a (a bloody/inflammatory strain) while the lowest was observed among strain A13a (a watery diarrhea associated strain; **Fig 6A**). Infection with bloody/inflammatory strains resulted in a 2-fold higher FITC-dextran translocation compared to watery strains (p = 0.0027; **Fig 6B**).

**Fig 6.**
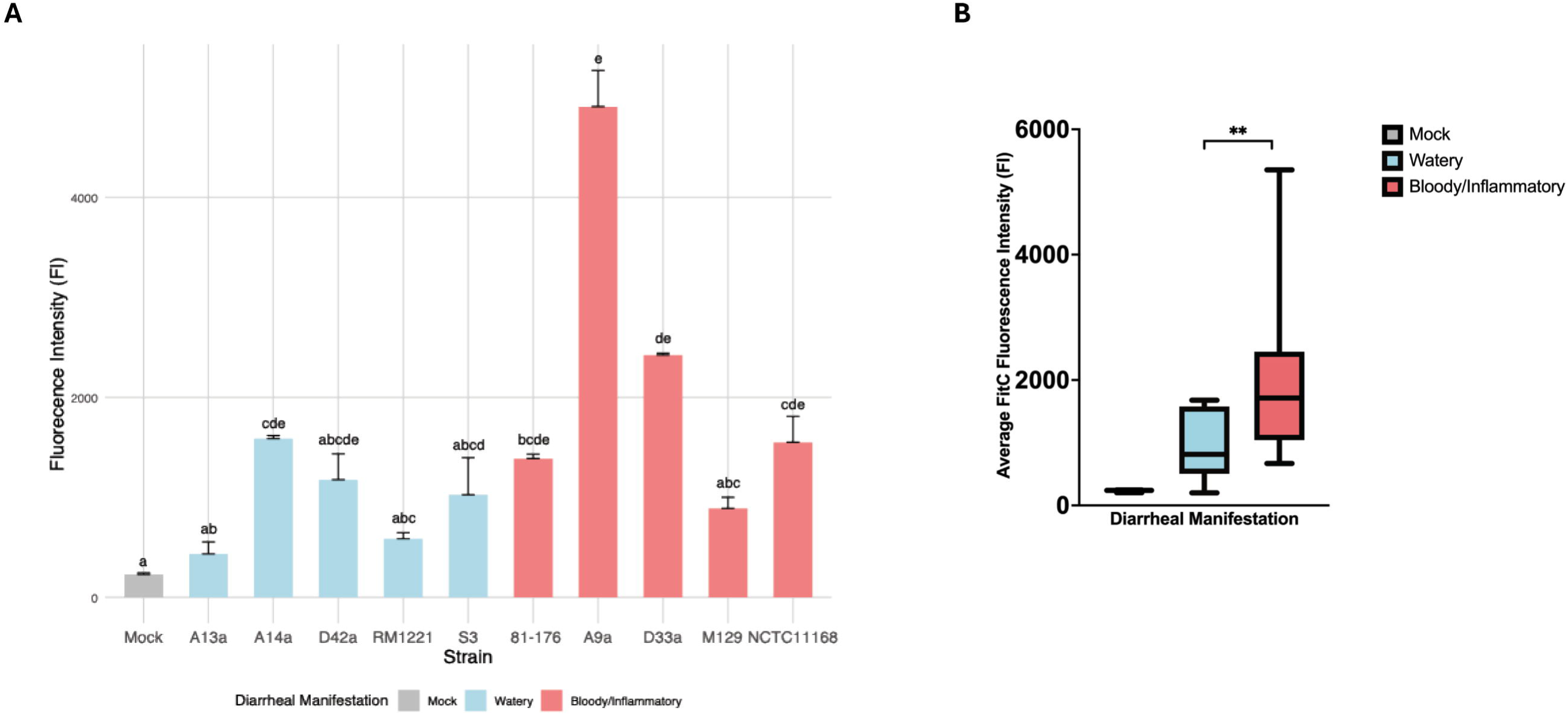
The fluorescence of FITC-Dextran flow through after 72 hours of infection with *C. jejuni* strains. (A) Average FITC-Dextran fluorescence intensity of each *C. jejuni* strain. Bars labeled with different letters indicate statistically significant differences (p < 0.05) based on Dunn’s post hoc test following a Kruskal-Wallis analysis (n = 3). Shared letters denote groups that are not significantly different from each other. (B) There was a highly significant difference between the watery and bloody/inflammatory diarrhea associated strains (**, p = 0.0027). Mock infection represents Caco-2 BBE cells that contained only sterile cell culture media.

### C. jejuni promotes translocation of commensal E. coli across the epithelial barrier

We next assessed whether barrier disruption by *C. jejuni* strains altered host susceptibility to microbial translocation. Using polarized Caco-2 BBE monolayers in Transwell inserts, we evaluated whether a non-invasive commensal strain of *E. coli* (DH5α) could cross the epithelial barrier during or after infection with *C. jejuni* strains. Two experimental approaches were tested: (i) co-inoculation of *C. jejuni* and *E. coli* for 6 hours, and (ii) pre-infection with *C. jejuni* for 72 hours followed by 3-hour exposure to *E. coli*. In both models, *E. coli* translocation was observed among all strains and there were no significant differences between strains (**Supplementary Fig S6A, S6C**). During co-inoculation, the mean *E. coli* CFU recovered from the basal chamber was not significantly different between bloody/inflammatory and watery diarrhea associated strains (p = 0.66; **Supplementary Fig S6B**). In the sequential infection model, basal recovery of *E. coli* was more pronounced, but no significant difference was observed between bloody/inflammatory and watery diarrhea associated strains (p = 0.30; **Supplementary Fig S6D**).

### Strains associated with bloody/inflammatory diarrhea demonstrate enhanced intracellular survival in macrophages

To investigate whether there was strain variation in the interactions with innate immune cells, we measured intracellular survival of the *C. jejuni* strains in J774A.1 murine macrophages at 24-, 48-, and 72-hours post-infection. There were significant differences between strains at all time points. Notably, strain A9a had the highest percent survivability across all time points while 81-176 had the lowest (**Fig 7A, 7C, 7E**). At 24 hours, viable intracellular *C. jejuni* was 2.6-fold higher among bloody/inflammatory diarrhea associated strains but there was no significant difference compared to watery diarrhea associated strains (p = 0.23; **Fig 7B**). At 48 hours post infection, bloody/inflammatory diarrhea associated strains showed significantly greater intracellular survivability (p = 0.011; **Fig 7D**). However, by 72-hours post infection, there was again no significant difference between bloody/inflammatory and watery diarrhea associated strains (p = 0.43; **Fig 7F**).

**Fig 7.**
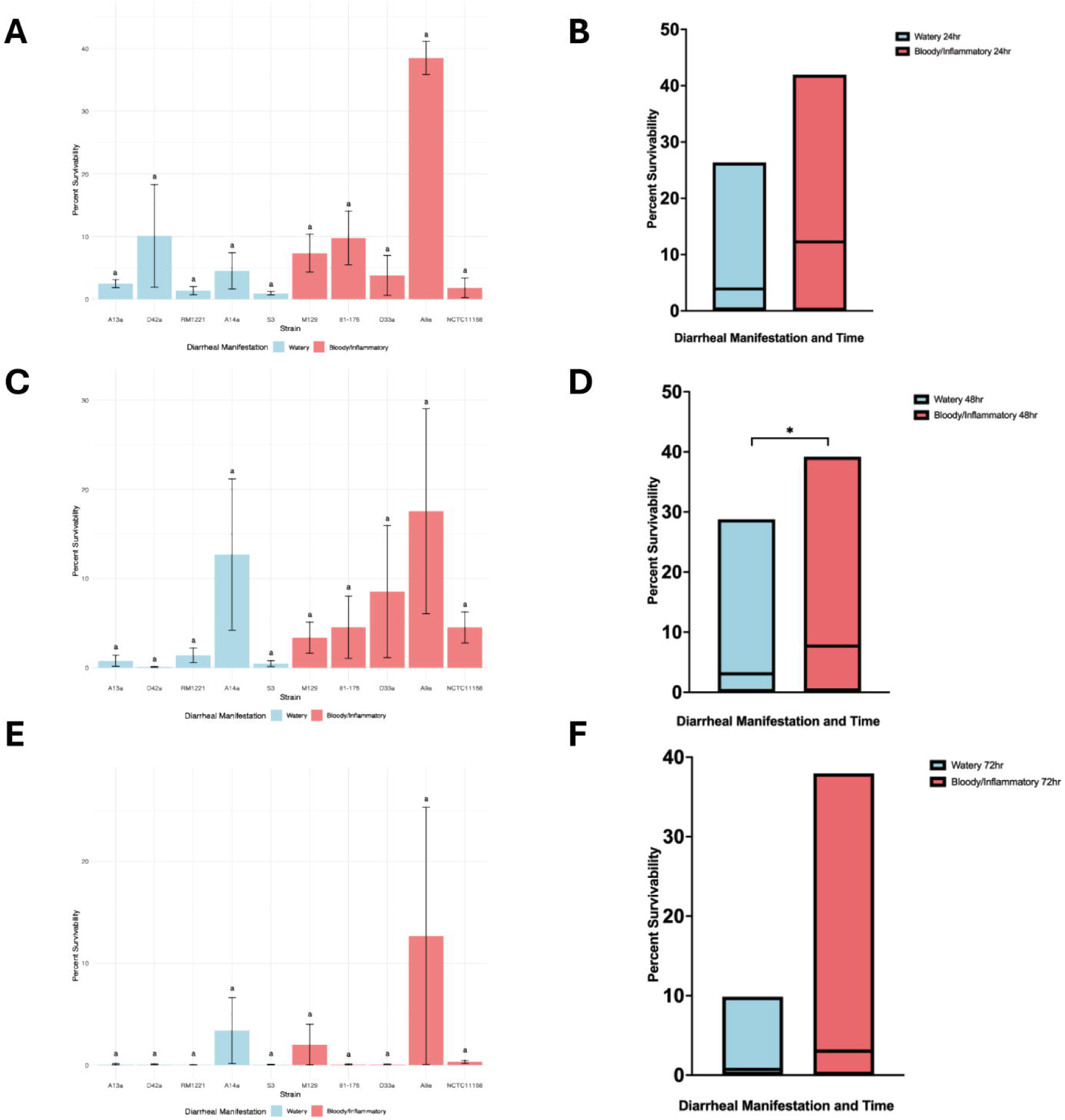
Survivability of *C. jejuni* strains in J774A.1 macrophages over 24, 48 and 72 hours. (A) Survivability by each *C. jejuni* strain over 24 hours. Bars labeled with different letters indicate statistically significant differences (p < 0.05) based on Dunn’s post hoc test following a Kruskal-Wallis analysis. Shared letters denote groups that are not significantly different from each other. Average percent survivability was calculated based on three biologically independent assays with two technical replicates (n = 3). (B) Macrophage survivability of *C. jejuni* strains grouped by diarrheal manifestation over 24 hours (p = 0.23). (C) Survivability by each *C. jejuni* strain over 48-hours. (D) Macrophage survivability of *C. jejuni* strains grouped by diarrheal manifestation over 48 hours (*, p = 0.011). (E) Survivability of *C. jejuni* strains over 72 hours. (F) Macrophage survivability of *C. jejuni* strains grouped by diarrheal manifestation over 72 hours (p = 0.43).

## Discussion

Clinical outcomes associated with different *C. jejuni* strains vary widely, from mild, self-limiting watery diarrhea to more severe presentations involving bloody and/or inflammatory diarrhea, abdominal cramping, and fever[10]. While host factors such as immune status and microbiota composition influence disease severity, accumulating evidence suggests that bacterial strain differences also contribute significantly to clinical heterogeneity[6, 32]. However, identifying phenotypic or molecular markers that distinguish high- and low-virulence *C. jejuni* strains has proven challenging[33]. While group-level differences were clear, we observed notable variation within each group, consistent with strain-to-strain variations from studies mentioned previously. Some watery strains exhibited moderate invasion or barrier disruption, and not all bloody/inflammatory strains displayed the most extreme phenotypes. A study by Law et al performed invasion, attachment and macrophage survivability assays using six out of the 10 *C. jejuni* strains used in this study[21]. In contrast to our study which used human intestinal epithelial cells (Caco-2 BBE, INT-407 and T84), the previous study used non-polarized human epithelial cells (HEp-2) that don’t form tight junctions for attachment and invasion assays. Our study has shown that different cell lines can provide very different results among *C. jejuni* strains. Furthermore, macrophage survivability was only performed at 72 hours post infection, missing significant differences at 48 hours that exist between watery and bloody/inflammatory diarrhea associated strains. This study dramatically expands from the Law et al study by examining a more detailed phenotypic analysis between *C. jejuni* strains, in addition to approaching the strains as two groups with distinct diarrheal manifestations confirmed in neonatal piglets.

Using a neonatal piglet model, which closely recapitulates human *C. jejuni* associated diarrheal disease[21, 29, 34], *C. jejuni* strains can be classified into two groups that consistently induce a particular diarrheal manifestation of either watery diarrhea or bloody/inflammatory in the neonatal piglet model[21, 34, 35]. The strains used in this study were chosen for their known diarrheal manifestation as well as their varied source background of either clinical or poultry sources. Given that poultry are the primary reservoirs of *C. jejuni*, our strains represent both mild and severe infections that can arise from poultry. Individuals experiencing milder, self-limiting symptoms such as watery diarrhea may be less likely to seek medical attention, leading to a lack of clinical isolates. By comparing *in vitro* virulence traits, we found significant strain to strain variation both within and between diarrheal manifestation groups, particularly among strains associated with bloody/inflammatory diarrhea which consistently exhibited enhanced invasion, intracellular persistence, epithelial barrier disruption and macrophage survivability, each a hallmark of a more invasive and inflammatory disease mechanism.

Adherence to intestinal epithelial cells is one of the primary steps to initiating *C. jejuni* infection[36, 37]. Here we found no significant differences in adherence levels between watery and bloody/inflammatory diarrhea associated strains to small intestinal INT407 cells or polarized large intestinal cells Caco-2 BBE and T84. Non-polarized cells such as INT407 cells lack a defined brush border and other features such as tight junctions that are more representative of the epithelial cells of the intestinal tract[38]. While as a group adherence levels were not significantly different, there was strain to strain variation observed, particularly with strain M129 having higher adherence levels than strains D33a, A9a, NCTC11168, and RM1221. Other studies have observed higher INT407 adherence among *C. jejuni* strains associated with febrile diarrhea or those from clinical sources compared to strains from patients without diarrhea/fever or from environmental sources[39, 40]. Similarly, adherence levels were not significantly different between groups when using Caco-2 BBE cells, a differentiated epithelial cell line originating from the human colon[41].

Using this cell line, there was lower strain to strain variation, where only 81-176 had significantly lower levels of adherence than strain A9a. Polarized cells such as Caco-2 BBE cells are able to form polarized monolayers with microvilli, and tight junctions, in addition to displaying brush border proteins that, despite being of colonic origin, are characteristic of small intestinal villus cells, including villin, fimbrin and the terminal web proteins fodrin and myosin II[42–45]. Adherence to T84 cells was similar across all *C. jejuni* strains and no significant differences were observed between diarrheal manifestation groups or between strains. T84 cells are derived from a human colonic carcinoma and are highly polarized with short microvilli, tight junctions and desmosomes between cells[46]. T84 cells offer a more accurate model of colonocytes due to their retention of key colonic features, including high expression of colonic markers and the presence of short microvilli characteristic of colonocytes[47]. Variation in the adherence of *C. jejuni* strains among polarized and non-polarized tissue culture cell lines requires more investigations into the mechanisms driving this individual strain variation. However, our findings support that both watery and bloody/inflammatory diarrhea associated strains are able to attach at similar levels to intestinal epithelial cells from the small and large intestine, however their invasiveness once at that epithelial surface is where they really differ.

Invasion has long been regarded as a hallmark of *C. jejuni* pathogenesis and has been studied extensively in tissue culture models[37, 48–51]. In the present study we found that bloody/inflammatory diarrhea associated strains demonstrated significantly higher invasiveness of both non-polarized and polarized intestinal epithelial cells. While the individual invasiveness of bloody/inflammatory diarrhea strains varied, there were few significant differences between strains from the same diarrheal manifestation, as all bloody/inflammatory diarrhea associated strains were invasive and all watery diarrhea associated strains were poor to non-invasive. Several groups have also observed variation in the invasiveness of *C. jejuni* strains. Everest et al (1992) found that *C. jejuni* clinical isolates from patients with colitis were more invasive than isolates from patients with non-inflammatory diarrhea[52]. Strain dependent invasiveness has also been observed using environmental strains compared to clinical strains, where clinical strains were found to be significantly more invasive[40]. Even intra-strain invasiveness has been found, particularly among continuously passed laboratory strains. For example, the clinical strain 81-176 has been reported to range in its invasiveness from 0.8 to 1.8%[53–55], while the genome sequenced clinical strain NCTC11168 (NCTC11168-GS) had diminished invasiveness and translocation abilities compared to the original isolate (NCTC11168-O), highlighting the potential effects of high laboratory passage on strain pathogenicity and colonization[56]. Furthermore, Cooper et al (2013) found that NCTC11168-GS had decreased disease production in the neonatal piglet model compared to NCTC11168-O[35]. Taken together, the consistent invasiveness observed among bloody/inflammatory diarrhea associated strains in our study, regardless of cell line used, supports invasion as a defining characteristic of the bloody/inflammatory diarrheal manifestation. The lack of invasiveness among watery diarrhea-associated strains may reflect alternative pathogenic strategies, such as growth capabilities and metabolic adaptability, that do not rely on epithelial invasion. These differences in invasive capacity likely contribute to the distinct clinical presentations observed between strain groups and reinforce the need for broader strain inclusion in *C. jejuni* pathogenesis research.

Bloody/inflammatory strains not only exhibit greater invasion of both polarized and non-polarized epithelial cells but also demonstrate enhanced intracellular persistence. Intracellular survival of *C. jejuni* has been linked to proinflammatory signaling and immune evasion in both *in vitro* and *in vivo* models [31, 57], but this is the first study to demonstrate that such phenotypes are heightened in strains associated with a consistent bloody/inflammatory diarrheal manifestation *in vivo*. Through macrophage survivability assays, we observed a connection to immune evasion by bloody/inflammatory diarrhea associated strains where these strains exhibited prolonged persistence within macrophages up to 48 hours post-infection. This suggests that not only are bloody/inflammatory diarrhea associated strains able to survive for at least 24 hours within intestinal epithelial cells, but they also have an ability to evade phagocytic killing for several days. Strain variation in macrophage survivability was most pronounced between bloody/inflammatory diarrhea strains, particularly among strain A9a at 24 hours where it had higher levels of survivability, although not statistically significant, compared to all other strains. Consistent with our study, Day et al (2000) and Law et al (2009) found that M129 was able to be recovered from J774A.1 macrophages after 72-hours, while the later study also observed survivability among A9a, A13a, A14a and D42a[21, 58]. Survivability differences between watery and bloody/inflammatory diarrhea associated strains at 48 hours, but not at 24 or 72 hours, suggests a transient but critical window during which certain *C. jejuni* strains, particularly those associated with bloody/inflammatory diarrhea, are better equipped or contain alternative mechanisms to persist slightly longer within host immune cells. *C. jejuni* contains one catalase enzyme, encoded by the *katA* gene, which aids in protection from reactive oxygen species generated during the oxidative burst once the bacterium is phagocytosed within macrophages[59, 60]. The *katA* gene has been found to be critical for *C. jejuni* macrophage survivability, where wildtype strain M129 was shown to be able to persist at levels significantly higher than a M129 *katA* mutant at 24, 48 and 72 hours within J774A.1 macrophages[58]. Another study found that the downregulation of *katA* in *C. jejuni* strain NCTC11168 through a mutation in a MarR-type transcriptional regulator (Cj1556) was also reduced in J774A.1 macrophage survivability[61, 62]. Future studies should investigate catalase expression or activity contributes to the prolonged intracellular persistence observed in bloody/inflammatory diarrhea-associated strains.

Disruption of epithelial barrier function further differentiated the two diarrheal manifestation groups. Bloody/inflammatory strains induced greater reductions in TEER and increased paracellular flux of FITC-dextran across polarized monolayers - both indicators of tight junction impairment[63, 64]. Other studies evaluating the changes to the TEER *in vitro* utilizing invasive *C. jejuni* strains, particularly 81-176 and NCTC11168, did find a significant drop in the TEER. However, none of these studies utilized strains with defined diarrheal manifestations or compared them to watery diarrhea associated strains[65–68]. This disruption in barrier integrity was significantly more pronounced in bloody/inflammatory strains and our study is the first to directly link this phenotype with reproducible diarrheal manifestations across multiple isolates. The effects we observed on the integrity of the Caco-2 BBE monolayers, regardless of defined diarrheal manifestation, translated into increased translocation of a non-invasive commensal *E. coli* strain, providing evidence that the barrier compromise facilitated microbial movement across the epithelium. The translocation of noninvasive bacteria such as *E. coli* during *C. jejuni* infection has been previously investigated *in vitro* with varying results. Konkel et al (1992) observed no change in the TEER or translocation of *E. coli* DH5α up to 6 hours following inoculation of Caco-2 cell monolayers using bloody/inflammatory strain M129 and strains of unknown diarrheal manifestations; F38011, 81116 and 78-27[69]. In our study, despite watery diarrhea associated strains being poorly invasive or non-invasive, we observed *E. coli* translocation at similar levels to bloody/inflammatory strains, suggesting alternate means for *E. coli* translocation other than invasion. Translocation of *E. coli* was found to not correspond with *C. jejuni* invasiveness in one study that used invasion-defective *C. jejuni* flagella mutant strains, with lipid raft-mediated endocytosis suggested to induce this effect[70]. The use of M cells to promote transcytosis of non-invasive bacteria was also observed using bloody/inflammatory diarrhea associated strain 81-176 but was not investigated using any other strains[71]. Both lipid rafts and M cells should be further investigated as potential mechanisms behind *E. coli* translocation by invasive and non-invasive *C. jejuni* strains.

Not all measured pathogenicity phenotypes differed between the two manifestation groups. Our study found that CDT production levels were not altered both between strains and across diarrheal manifestation groups at either temperature, suggesting CDT is not a discriminating factor for diarrheal severity. CDT is widespread among *C. jejuni* isolates and contributes to host cell cycle arrest, DNA damage, and epithelial disruption[72], however the specific contributions of CDT in *C. jejuni* pathogenesis remains unclear[73, 74]. There is limited evidence that CDT expression is critical to disease but it was found that CDT contributes to greater adherence and invasion of epithelial cells along with more progressive gastritis and proximal duodenitis in two *in vivo* mouse models[75, 76]. However, the role of CDT in driving different diarrheal symptoms is not well understood. Previous studies have assessed CDT production with HeLa cells among *C. jejuni* strains from clinical or poultry sources and have reported variability, although none had a diarrheal manifestation associated with any of the strains[77–79]. Additionally, IL-8 secretion, a hallmark of initial epithelial response to *C. jejuni* contact[80–82] and a potent chemoattractant and cellular activator[81] also did not differ significantly between diarrheal manifestation groups at 24 hours post-infection. Although not statistically significant, strain to strain variation was observed among bloody/inflammatory strains with D33a and NCTC11168 having lower IL-8 levels than M129 and A9a. Watery diarrhea associated strains showed less variation between strains with only A13a and A14a having lower IL-8 levels than other watery diarrhea associated strains. Few studies have looked at differences in IL-8 secretion across *C. jejuni* strains and this is the first study to assess IL-8 secretion by *C. jejuni* strains with known diarrheal manifestations in the neonatal piglet model. One study using bloody/inflammatory diarrhea associated strain NCTC11168 and human colitis strain L115 showed no difference in IL-8 secretion by T84 cells[83], while another study looked at IL-8 secretion between *C. jejuni* strains from poultry and human sources and found no difference in IL-8 secretion by T84 cells between groups[84]. However, IL-8 secretion is not based solely on the invasive or adherent capability of *C. jejuni* strains, as it can also be activated by CDT production[81, 82], indicating that early chemokine responses may not correlate with longer-term invasive or inflammatory phenotypes. Furthermore, our findings that both bloody/inflammatory and watery diarrhea associated strains produce similar levels of CDT and elicit comparable IL-8 responses despite significant differences in invasion. This variation highlights the importance of considering strain-level differences when evaluating the inflammatory potential of *C. jejuni* and suggests that multiple mechanisms may drive IL-8 secretion in the host.

Strain A9a, originally recovered from a poultry processing plant in Kansas, USA[85], is associated with bloody/inflammatory diarrhea in the neonatal piglet model. A9a demonstrated the highest levels of adherence, invasion, and intracellular survival to intestinal epithelial cells compared to all other *C. jejuni* strains. While its CDT production was comparable to many of the other *C. jejuni* strains, A9a showed the highest stimulation of IL-8 secretion from T84 cells, results that support IL-8 production due to high adherence and invasion even with similar levels of CDT. A9a also caused the most pronounced disruption to epithelial barrier integrity, with the steepest decline in TEER, the highest FITC-dextran permeability, and the greatest translocation of *E. coli* DH5α across Caco-2 BBE monolayers over 72 hours. Finally, A9a had the highest survivability within macrophages across all time points, reflecting its resilience against innate immune responses. Despite its strong *in vitro* virulence potential, A9a was not hypervirulent in the piglet model[21]. Instead, it produced bloody/inflammatory diarrhea at similar levels to the other strains in this study, 81-176, M129, D33a and NCTC11168, suggesting that multiple factors may drive the severity of diarrhea. *C. jejuni* strain variation has been observed in other studies, such as Ex114, a *C. jejuni* strain that was found to be 25 times more invasive than reference strain 81116, which is a relatively low invader[16]. Consistent with our findings, the same study reported that increased invasiveness was not linked to enhanced adhesion, and motility levels were similar between highly and minimally invasive strains. In other studies, stain 81-176, sourced from a raw milk outbreak,[86] is widely regarded as a highly invasive clinical isolate and has been shown to cause robust epithelial invasion in cell lines[53, 87] and animal models[88] as well as produce disease in humans[89]. Yet its performance varies between laboratories depending on passage history[53, 55, 90]. Repeatedly passages reduced the invasiveness and serum survival of 81-176[91]. Similarly, while strain NCTC11168 is a reference strain in *C. jejuni* research, its virulence is known to diminish with laboratory passage, leading to reduced motility, invasion and colonization capacity *in vitro*[56] in addition to altering *in vivo* virulence in the neonatal piglet model[35]. Our findings support these previous reports that strain-dependent variation plays a key role in pathogenicity.

This strain-to-strain variations observed in this study, even among strains associated with a particular diarrheal manifestation in the neonatal piglet model, highlight the inherent complexity and plasticity of *C. jejuni* virulence[92–95]. This study also underscores the need for population-scale studies integrating phenotypic data with comparative genomics to identify combinations of traits that drive disease severity. One such study that performed a comparative genomic analysis on 67 *C. jejuni* and 12 *C. coli* strains, identified the *Campylobacter jejuni*-integrated elements (CJIEs) originally identified in *C. jejuni* strain RM1221, were variably present across other *Campylobacter* strains[96]. These genomic elements, some of which included prophages and mobile genetic elements, contribute to functional differences that may affect virulence, stress response, or host interaction. Notably, Parker et al reported that coding sequences within CJIEs were highly variable, highlighting a potential genomic mechanism for the phenotypic diversity observed in our assays[96]. This supports our findings that even within clinically defined groups, virulence phenotypes can differ, potentially due to underlying genomic heterogeneity. One limitation of this study is that the number of strains included was limited, but they were well-characterized and associated with consistent clinical phenotypes in a validated *in vivo* neonatal piglet model. Our *in vitro* assays cannot fully replicate host complexity, but the consistency of results across multiple models strengthens confidence in the observed trends. Future work should integrate whole-genome sequencing, transcriptomic profiling, and *in vivo* validation to identify the molecular determinants responsible for these virulence phenotypes.

In summary, *C. jejuni* strains associated with bloody/inflammatory diarrhea exhibit a distinct virulence profile characterized by epithelial invasion, intracellular survival, barrier disruption, and macrophage persistence (**Fig 8**). Together, enhanced invasion and intestinal damage, intracellular survival, and macrophage survivability provide a functional explanation for the more severe pathology observed in the piglet model by bloody/inflammatory strain and, by extension, potentially in human infections. By linking *in vivo* disease phenotypes to mechanistic *in vitro* traits, our findings advance understanding of *C. jejuni* pathogenesis and provide a framework for developing targeted diagnostics, risk-based screening tools, and novel therapeutic interventions. Furthermore, the strain-to-strain variation observed in this study, even within groups associated with the same *in vivo* diarrheal manifestation, present that caution is needed when interpreting the results of experiments that utilize only a few *C. jejuni* strains.

**Fig 8.**
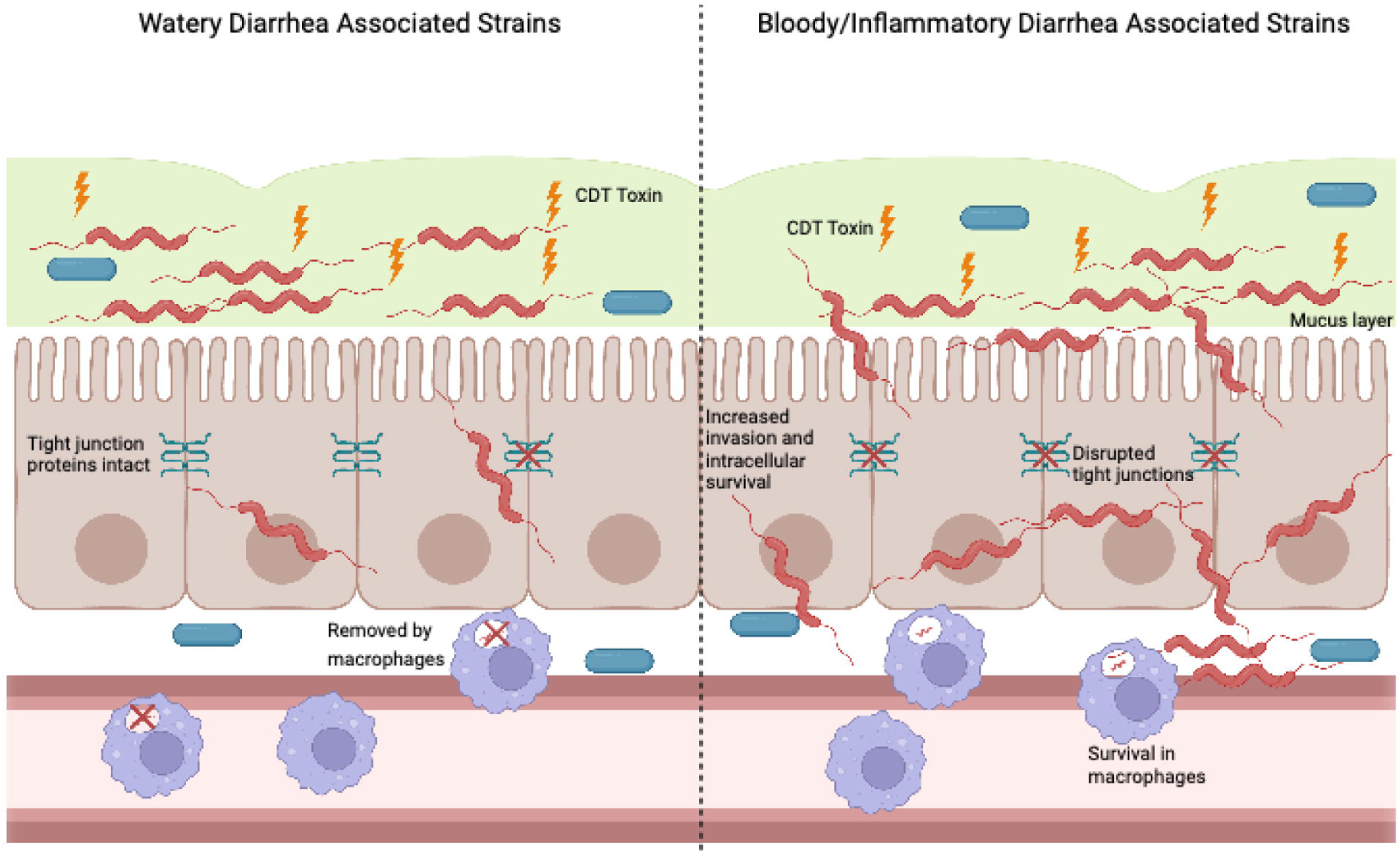
Proposed lifestyles of *C. jejuni* strains associated with watery or bloody/inflammatory diarrhea. Watery diarrhea associated strains live an extracellular lifestyle where they attach to enterocytes and produce CDT toxin. They are poor invaders, disrupting the tight junctions between epithelial cells to a minimal degree, although commensal bacterial such as *E. coli* are translocated. Those that do weakly invade, do not survive long term intracellularly and are effectively removed by macrophages. Bloody/inflammatory diarrhea associated strains support an intracellular lifestyle where they attach to and invade enterocytes, significantly disrupting the tight junctions between those enterocytes. Bloody/inflammatory diarrhea associated strains can evade killing by macrophages for longer periods of time, promoting infection and contributing to more severe patient symptoms. Figure created in BioRender.

## Acknowledgments

The authors thank all members of the Cooper laboratory for providing critical technical assistance in sample processing and other aspects of the study.

## Ethics statement

This study did not involve the use of animals or human participants. The *Campylobacter jejuni* strains used in this study were previously characterized in a neonatal piglet model under approved animal use protocols, as described in Law et al and Cooper et al. No new *in vivo* experiments were conducted for this study.

## Data availability statement

All relevant data are within the manuscript and its supplementary files.

## Author Contributions

**JB:** Conceptualization, methodology, experimentation, formal analysis, writing – original draft, and review & editing. **KKC:** Conceptualization, methodology, resources, funding acquisition, project administration, supervision, writing, and review & editing. **CP:** Writing, and review & editing. **BP:** Writing, and review & editing.

## Funding statement

This work was supported in part by Start-up funds provided to Kerry Cooper by the University of Arizona, and Technology and Research Initiative Fund (TRIF) award CALS_ACBC_Cooper_2101696 provided to Kerry Cooper by the University of Arizona. No funding agency had any role in the study design, data collection, data analysis, or preparation of the manuscript.

## Competing interests

The authors declare no competing interests.

## Supplementary Figure Legends

**Supplementary Fig S1. Invasion of INT407 cells.** (A) Invasion across all ten *C. jejuni* strains. Bars labeled with different letters indicate statistically significant differences (p < 0.05) based on Dunn’s post hoc test following a Kruskal-Wallis analysis. Shared letters denote strains that are not significantly different from each other. (B) Invasiveness of INT407 cells by *C. jejuni* strains grouped by diarrheal manifestation (***, p = 0.0006).

**Supplementary Fig S2. Average percent invasion of intestinal epithelial cell line T84.** (A) Invasion of T84 cells across all ten *C. jejuni* strains. Bars labeled with different letters indicate statistically significant differences (p < 0.05) based on Dunn’s post hoc test following a Kruskal-Wallis analysis. Shared letters denote groups that are not significantly different from each other. (B) Invasiveness of *C. jejuni* strains grouped by diarrheal manifestation (**, p = 0.0063).

**Supplementary Fig S3. Adherence to T84 cells.** (A) Adherence to T84 cells by *C. jejuni* strains. Bars labeled with different letters indicate statistically significant differences (p < 0.05) based on Dunn’s post hoc test following a Kruskal-Wallis analysis. Shared letters denote groups that are not significantly different from each other (n = 3). Strain 81-176 was not included in this data set. (B) Adherence to T84 cells by *C. jejuni* strains grouped by diarrheal manifestation (p = 0.90). Strain 81-176 is not included in this data set.

**Supplementary Fig S4. Cytolethal Distending Toxin (CDT) production at 37°C and 42°C**.

(A) CDT toxin production at 37°C and 42°C by *C. jejuni* strains. (B) CDT production *C. jejuni* strains grouped by diarrheal manifestation at 37°C (p = 0.74) and 42°C (p = 0.086). CDT production was not significantly different when performed at 37°C compared to 42°C for watery diarrhea associated strains (p = 0.65) or bloody/inflammatory diarrhea associated strains (p = 0.12).

**Supplementary Fig S5. IL-8 Production following 24-hour incubation with *C. jejuni* strains**.

(A) IL-8 production in pg/ml of *C. jejuni* strains. Bars labeled with different letters indicate statistically significant differences (p < 0.05) based on Dunn’s post hoc test following a Kruskal-Wallis analysis. Shared letters denote groups that are not significantly different from each other. Strain 81-176 is not included in this data set. Mock infection represents Caco-2 BBE cells that contained only sterile cell culture media. (B) IL-8 production in pg/ml of *C. jejuni* strains grouped by diarrheal manifestation (p = 0.61). Mock infection represents Caco-2 BBE cells that contained only sterile cell culture media. Strain 81-176 is not included in this data set.

**Supplementary Fig S6. Translocation of *Escherichia coli* through Caco-2 BBE cell monolayers.** (A) 6-hour co-culture incubation with *C. jejuni* strains. Bars labeled with different letters indicate statistically significant differences (p < 0.05) based on Dunn’s post hoc test following a Kruskal-Wallis analysis. Shared letters denote groups that are not significantly different from each other. (B) 6-hour co-culture incubation with *C. jejuni* strains grouped by diarrheal manifestation (p = 0.66). Mock infection represents Caco-2 BBE cells that contained only sterile cell culture media. (C) Pre-infection with *C. jejuni* for 72 hours followed by 3-hour exposure to *E. coli* through Caco-2 BBE cells by strain (n = 3). (D) Pre-infection with *C. jejuni* for 72 hours followed by 3-hour exposure to *E. coli* through Caco-2 BBE grouped by diarrheal manifestation (p = 0.30).

## Notes

### Competing Interest Statement

The authors have declared no competing interest.

### Summary of Updates

This version of the manuscript has been revised to correct an error in the statistical analysis, which has resulted in the epithelial cell attachment no longer being statistically significant. All figures have been revised to reflect the new statistically analysis.

## References

1. Organization WH. Global View of Campylobaceriosis, Report of an Expert Consultation. 2012.

2. Walter EJS, Cui Z, Tierney R, Griffin PM, Hoekstra RM, Payne DC, et al. Foodborne Illness Acquired in the United States—Major Pathogens, 2019. Volume 31, Number 4. 2025. PubMed PMID: April.

3. Schorling E, Knorr S, Lick S, Steinberg P, Brüggemann DA. Probability of sequelae following *Campylobacter* spp. infections: Update of systematic reviews and meta-analyses. Public Health Challenges. 2023;2(4):e145. doi: 10.1002/puh2.145.

4. Nyati KK, Nyati R. Role of *Campylobacter jejuni* infection in the pathogenesis of Guillain-Barré syndrome: an update. Biomed Res Int. 2013;2013:852195. Epub 20130813. doi: 10.1155/2013/852195. PubMed PMID: 24000328; PubMed Central PMCID: PMCPMC3755430.

5. Peters S, Pascoe B, Wu Z, Bayliss SC, Zeng X, Edwinson A, et al. *Campylobacter jejuni* genotypes are associated with post-infection irritable bowel syndrome in humans. Commun Biol. 2021;4(1):1015. Epub 20210830. doi: 10.1038/s42003-021-02554-8. PubMed PMID: 34462533; PubMed Central PMCID: PMCPMC8405632.

6. Kotloff KL, Nataro JP, Blackwelder WC, Nasrin D, Farag TH, Panchalingam S, et al. Burden and aetiology of diarrhoeal disease in infants and young children in developing countries (the Global Enteric Multicenter Study, GEMS): a prospective, case-control study. Lancet. 2013;382(9888):209-22. Epub 20130514. doi: 10.1016/s0140-6736(13)60844-2. PubMed PMID: 23680352.

7. Investigators TM-EN. The MAL-ED Study: A Multinational and Multidisciplinary Approach to Understand the Relationship Between Enteric Pathogens, Malnutrition, Gut Physiology, Physical Growth, Cognitive Development, and Immune Responses in Infants and Children Up to 2 Years of Age in Resource-Poor Environments. Clinical Infectious Diseases. 2014;59(suppl_4):S193–S206. doi: 10.1093/cid/ciu653.

8. Haque MA, Platts-Mills JA, Mduma E, Bodhidatta L, Bessong P, Shakoor S, et al. Determinants of *Campylobacter* infection and association with growth and enteric inflammation in children under 2 years of age in low-resource settings. Sci Rep. 2019;9(1):17124. Epub 20191120. doi: 10.1038/s41598-019-53533-3. PubMed PMID: 31748573; PubMed Central PMCID: PMCPMC6868199.

9. Pogreba-Brown K, Baker A, Ernst K, Stewart J, Harris RB, Weiss J. Assessing risk factors of sporadic *Campylobacter* infection: a case-control study in Arizona. Epidemiol Infect. 2016;144(4):829–39. doi: 10.1017/s0950268815002162. PubMed PMID: 26828241.

10. Igwaran A, Okoh AI. Human campylobacteriosis: A public health concern of global importance. Heliyon. 2019;5(11):e02814. Epub 20191114. doi: 10.1016/j.heliyon.2019.e02814. PubMed PMID: 31763476; PubMed Central PMCID: PMCPMC6861584.

11. Harvey P, Battle T, Leach S. Different invasion phenotypes of *Campylobacter* isolates in Caco-2 cell monolayers. J Med Microbiol. 1999;48(5):461–9. doi: 10.1099/00222615-48-5-461. PubMed PMID: 10229543.

12. Ruiz-Palacios G, Torres N, Ruiz-Palacios B, Torres J, Escamilla E, Tamayo J. Cholera-like enterotoxin produced by *Campylobacter jejuni*: characterisation and clinical significance. The Lancet. 1983;322(8344):250–3.

13. Johnson WM, Lior H. Toxins produced by *Campylobacter jejuni* and *Campylobacter coli*. Lancet. 1984;1(8370):229-30. doi: 10.1016/s0140-6736(84)92155-x. PubMed PMID: 6141372.

14. Apel D, Ellermeier J, Pryjma M, Dirita VJ, Gaynor EC. Characterization of *Campylobacter jejuni* RacRS reveals roles in the heat shock response, motility, and maintenance of cell length homogeneity. J Bacteriol. 2012;194(9):2342–54. Epub 20120217. doi: 10.1128/JB.06041-11. PubMed PMID: 22343300; PubMed Central PMCID: PMCPMC3347078.

15. Turonova H, Briandet R, Rodrigues R, Hernould M, Hayek N, Stintzi A, et al. Biofilm spatial organization by the emerging pathogen *Campylobacter jejuni*: comparison between NCTC 11168 and 81-176 strains under microaerobic and oxygen-enriched conditions. Front Microbiol. 2015;6:709. Epub 20150713. doi: 10.3389/fmicb.2015.00709. PubMed PMID: 26217332; PubMed Central PMCID: PMCPMC4499754.

16. Fearnley C, Manning G, Bagnall M, Javed MA, Wassenaar TM, Newell DG. Identification of hyperinvasive *Campylobacter jejuni* strains isolated from poultry and human clinical sources. J Med Microbiol. 2008;57(Pt 5):570–80. doi: 10.1099/jmm.0.47803-0. PubMed PMID: 18436589.

17. Hofreuter D, Novik V,, Galán JE, . Metabolic Diversity in Campylobacter jejuni Enhances Specific Tissue Colonization. 2008.

18. Poly F, Ewing C, Goon S, Hickey TE, Rockabrand D, Majam G, et al. Heterogeneity of a *Campylobacter jejuni* protein that is secreted through the flagellar filament. Infect Immun. 2007;75(8):3859–67. Epub 20070521. doi: 10.1128/IAI.00159-07. PubMed PMID: 17517862; PubMed Central PMCID: PMCPMC1951984.

19. Meurens F, Summerfield A, Nauwynck H, Saif L, Gerdts V. The pig: a model for human infectious diseases. Trends Microbiol. 2012;20(1):50–7. Epub 20111205. doi: 10.1016/j.tim.2011.11.002. PubMed PMID: 22153753; PubMed Central PMCID: PMCPMC7173122.

20. Boosinger TR, Powe TA. *Campylobacter jejuni* infections in gnotobiotic pigs. Am J Vet Res. 1988;49(4):456–8. PubMed PMID: 3377304.

21. Law BF, Adriance SM, Joens LA. Comparison of in vitro virulence factors of *Campylobacter jejuni* to in vivo lesion production. Foodborne Pathog Dis. 2009;6(3):377–85. doi: 10.1089/fpd.2008.0183. PubMed PMID: 19278341.

22. Canibe N, O’Dea M, Abraham S. Potential relevance of pig gut content transplantation for production and research. Journal of Animal Science and Biotechnology. 2019;10(1):55. doi: 10.1186/s40104-019-0363-4.

23. van der Laan JW, Brightwell J, McAnulty P, Ratky J, Stark C. Regulatory acceptability of the minipig in the development of pharmaceuticals, chemicals and other products. J Pharmacol Toxicol Methods. 2010;62(3):184–95. Epub 20100616. doi: 10.1016/j.vascn.2010.05.005. PubMed PMID: 20601024.

24. Lodemann U, Amasheh S, Radloff J, Kern M, Bethe A, Wieler LH, et al. Effects of Ex Vivo Infection with ETEC on Jejunal Barrier Properties and Cytokine Expression in Probiotic-Supplemented Pigs. Dig Dis Sci. 2017;62(4):922–33. Epub 20161219. doi: 10.1007/s10620-016-4413-x. PubMed PMID: 27995406.

25. Gefeller EM, Martens H, Aschenbach JR, Klingspor S, Twardziok S, Wrede P, et al. Effects of age and zinc supplementation on transport properties in the jejunum of piglets. J Anim Physiol Anim Nutr (Berl). 2015;99(3):542–52. Epub 20140721. doi: 10.1111/jpn.12232. PubMed PMID: 25039419.

26. Lodemann U, Dillenseger A, Aschenbach JR, Martens H. Effects of age and controlled oral dosing of Enterococcus faecium on epithelial properties in the piglet small intestine. Benef Microbes. 2013;4(4):335–44. doi: 10.3920/bm2013.0004. PubMed PMID: 24311317.

27. Harvey RB, Young CR, Ziprin RL, Hume ME, Genovese KJ, Anderson RC, et al. Prevalence of *Campylobacter* spp isolated from the intestinal tract of pigs raised in an integrated swine production system. J Am Vet Med Assoc. 1999;215(11):1601–4. PubMed PMID: 14567422.

28. Taylor D, Olubunmi P. A re-examination of the role of Campylobacter fetus subspecies coli in enteric disease of the pig. The Veterinary Record. 1981;109(6):112–5.

29. Vítovec J, Koudela B, Štěrba J, Tomancová I, Matyáš Z, Vladík P. The gnotobiotic piglet as a model for the pathogenesis of *Campylobacter jejuni* infection. Zentralblatt für Bakteriologie. 1989;271(1):91–103. doi: 10.1016/s0934-8840(89)80058-1.

30. Abuoun M, Manning G, Cawthraw SA, Ridley A, Ahmed IH, Wassenaar TM, et al. Cytolethal distending toxin (CDT)-negative *Campylobacter jejuni* strains and anti-CDT neutralizing antibodies are induced during human infection but not during colonization in chickens. Infect Immun. 2005;73(5):3053–62. doi: 10.1128/IAI.73.5.3053-3062.2005. PubMed PMID: 15845513; PubMed Central PMCID: PMCPMC1087314.

31. Watson RO, Galán JE. *Campylobacter jejuni* survives within epithelial cells by avoiding delivery to lysosomes. PLoS Pathog. 2008;4(1):e14. doi: 10.1371/journal.ppat.0040014. PubMed PMID: 18225954; PubMed Central PMCID: PMCPMC2323279.

32. Stamps BW, Kuroiwa J, Isidean SD, Schilling MA, Harro C, Talaat KR, et al. Exploring Changes in the Host Gut Microbiota During a Controlled Human Infection Model for *Campylobacter jejuni*. Frontiers in Cellular and Infection Microbiology. 2021;11. doi: 10.3389/fcimb.2021.702047.

33. Cody AJ, McCarthy NM, Wimalarathna HL, Colles FM, Clark L, Bowler IC, et al. A longitudinal 6-year study of the molecular epidemiology of clinical *Campylobacter* isolates in Oxfordshire, United kingdom. J Clin Microbiol. 2012;50(10):3193–201. Epub 20120718. doi: 10.1128/JCM.01086-12. PubMed PMID: 22814466; PubMed Central PMCID: PMCPMC3457434.

34. Babakhani FK, Bradley GA, Joens LA. Newborn piglet model for campylobacteriosis. Infect Immun. 1993;61(8):3466–75. doi: 10.1128/iai.61.8.3466-3475.1993. PubMed PMID: 8335377; PubMed Central PMCID: PMCPMC281024.

35. Cooper KK, Cooper MA, Zuccolo A, Joens LA. Re-sequencing of a virulent strain of *Campylobacter jejuni* NCTC11168 reveals potential virulence factors. Res Microbiol. 2013;164(1):6–11. Epub 20121006. doi: 10.1016/j.resmic.2012.10.002. PubMed PMID: 23046762.

36. Rubinchik S, Seddon A, Karlyshev AV. Molecular mechanisms and biological role of *Campylobacter jejuni* attachment to host cells. Eur J Microbiol Immunol (Bp). 2012;2(1):32–40. Epub 20120317. doi: 10.1556/EuJMI.2.2012.1.6. PubMed PMID: 24611119; PubMed Central PMCID: PMCPMC3933988.

37. Monteville MR, Yoon JE, Konkel ME. Maximal adherence and invasion of INT 407 cells by *Campylobacter jejuni* requires the CadF outer-membrane protein and microfilament reorganization. Microbiology (Reading). 2003;149(Pt 1):153–65. doi: 10.1099/mic.0.25820-0. PubMed PMID: 12576589.

38. Konkel ME, Hayes SF, Joens LA, Cieplak W, Jr. Characteristics of the internalization and intracellular survival of *Campylobacter jejuni* in human epithelial cell cultures. Microb Pathog. 1992;13(5):357–70. doi: 10.1016/0882-4010(92)90079-4. PubMed PMID: 1297914.

39. Fauchere JL, Rosenau A, Veron M, Moyen EN, Richard S, Pfister A. Association with HeLa cells of *Campylobacter jejuni* and *Campylobacter coli* isolated from human feces. Infect Immun. 1986;54(2):283–7. doi: 10.1128/iai.54.2.283-287.1986. PubMed PMID: 3770943; PubMed Central PMCID: PMCPMC260156.

40. Newell DG, McBride H, Saunders F, Dehele Y, Pearson AD. The virulence of clinical and environmental isolates of *Campylobacter jejuni*. J Hyg (Lond). 1985;94(1):45–54. doi: 10.1017/s0022172400061118. PubMed PMID: 3973380; PubMed Central PMCID: PMCPMC2129391.

41. Peterson MD, Mooseker MS. Characterization of the enterocyte-like brush border cytoskeleton of the C2BBe clones of the human intestinal cell line, Caco-2. J Cell Sci. 1992;102 (Pt 3):581-600. doi: 10.1242/jcs.102.3.581. PubMed PMID: 1506435.

42. Hughson EJ, Cutler DF, Hopkins CR. Basolateral secretion of kappa light chain in the polarised epithelial cell line, Caco-2. J Cell Sci. 1989;94 (Pt 2):327-32. doi: 10.1242/jcs.94.2.327. PubMed PMID: 2516089.

43. Lea T. Caco-2 Cell Line. In: Verhoeckx K, Cotter P, López-Expósito I, Kleiveland C, Lea T, Mackie A, et al., editors. The Impact of Food Bioactives on Health: in vitro and ex vivo models. Cham (CH): Springer Copyright 2015, The Author(s). 2015. p. 103–11.

44. Sambuy Y, De Angelis I, Ranaldi G, Scarino M, Stammati A, Zucco F. The Caco-2 cell line as a model of the intestinal barrier: influence of cell and culture-related factors on Caco-2 cell functional characteristics. Cell biology and toxicology. 2005;21:1–26.

45. Peterson MD, Bement WM, Mooseker MS. An in vitro model for the analysis of intestinal brush border assembly. II. Changes in expression and localization of brush border proteins during cell contact-induced brush border assembly in Caco-2BBe cells. J Cell Sci. 1993;105 (Pt 2):461–72. doi: 10.1242/jcs.105.2.461. PubMed PMID: 8408277.

46. Dharmsathaphorn K, McRoberts JA, Mandel KG, Tisdale LD, Masui H. A human colonic tumor cell line that maintains vectorial electrolyte transport. Am J Physiol. 1984;246(2 Pt 1):G204–8. doi: 10.1152/ajpgi.1984.246.2.G204. PubMed PMID: 6141741.

47. Devriese S, Van den Bossche L, Van Welden S, Holvoet T, Pinheiro I, Hindryckx P, et al. T84 monolayers are superior to Caco-2 as a model system of colonocytes. Histochemistry and Cell Biology. 2017;148(1):85–93. doi: 10.1007/s00418-017-1539-7.

48. Konkel ME, Joens LA. Adhesion to and invasion of HEp-2 cells by *Campylobacter* spp. Infect Immun. 1989;57(10):2984–90. doi: 10.1128/iai.57.10.2984-2990.1989. PubMed PMID: 2550368; PubMed Central PMCID: PMCPMC260759.

49. Rivera-Amill V, Kim BJ, Seshu J, Konkel ME. Secretion of the virulence-associated *Campylobacter* invasion antigens from *Campylobacter jejuni* requires a stimulatory signal. J Infect Dis. 2001;183(11):1607–16. Epub 20010501. doi: 10.1086/320704. PubMed PMID: 11343209.

50. Russell RG, Blake DC, Jr. Cell association and invasion of Caco-2 cells by *Campylobacter jejuni*. Infect Immun. 1994;62(9):3773–9. doi: 10.1128/iai.62.9.3773-3779.1994. PubMed PMID: 8063393; PubMed Central PMCID: PMCPMC303030.

51. Konkel ME, Talukdar PK, Negretti NM, Klappenbach CM. Taking Control: Campylobacter jejuni Binding to Fibronectin Sets the Stage for Cellular Adherence and Invasion. Frontiers in Microbiology. 2020;11. doi: 10.3389/fmicb.2020.00564.

52. Everest PH, Goossens H, Butzler JP, Lloyd D, Knutton S, Ketley JM, et al. Differentiated Caco-2 cells as a model for enteric invasion by *Campylobacter jejuni* and *C. coli*. J Med Microbiol. 1992;37(5):319–25. doi: 10.1099/00222615-37-5-319. PubMed PMID: 1433253.

53. Yao R, Burr DH, Doig P, Trust TJ, Niu H, Guerry P. Isolation of motile and non-motile insertional mutants of *Campylobacter jejuni*: the role of motility in adherence and invasion of eukaryotic cells. Mol Microbiol. 1994;14(5):883–93. doi: 10.1111/j.1365-2958.1994.tb01324.x. PubMed PMID: 7715450.

54. Doig P, Yao R, Burr DH, Guerry P, Trust TJ. An environmentally regulated pilus-like appendage involved in *Campylobacter* pathogenesis. Molecular Microbiology. 1996;20(4):885–94.

55. Yao R, Burr DH, Guerry P. CheY-mediated modulation of *Campylobacter jejuni* virulence. Mol Microbiol. 1997;23(5):1021–31. doi: 10.1046/j.1365-2958.1997.2861650.x. PubMed PMID: 9076738.

56. Gaynor Erin C, Cawthraw S, Manning G, MacKichan Joanna K, Falkow S, Newell Diane G. The Genome-Sequenced Variant of *Campylobacter jejuni* NCTC 11168 and the Original Clonal Clinical Isolate Differ Markedly in Colonization, Gene Expression, and Virulence-Associated Phenotypes. Journal of Bacteriology. 2004;186(2):503–17. doi: 10.1128/jb.186.2.503-517.2004.

57. Novik V, Hofreuter D, Galán JE. Identification of *Campylobacter jejuni* genes involved in its interaction with epithelial cells. Infect Immun. 2010;78(8):3540–53. Epub 20100601. doi: 10.1128/iai.00109-10. PubMed PMID: 20515930; PubMed Central PMCID: PMCPMC2916286.

58. Day WA, Jr., Sajecki JL, Pitts TM, Joens LA. Role of catalase in *Campylobacter jejuni* intracellular survival. Infect Immun. 2000;68(11):6337–45. doi: 10.1128/iai.68.11.6337-6345.2000. PubMed PMID: 11035743; PubMed Central PMCID: PMCPMC97717.

59. De Melo MA, Gabbiani G, Pechère JC. Cellular events and intracellular survival of *Campylobacter jejuni* during infection of HEp-2 cells. Infect Immun. 1989;57(7):2214–22. doi: 10.1128/iai.57.7.2214-2222.1989. PubMed PMID: 2731988; PubMed Central PMCID: PMCPMC313863.

60. Kiehlbauch JA, Albach RA, Baum LL, Chang KP. Phagocytosis of *Campylobacter jejuni* and its intracellular survival in mononuclear phagocytes. Infect Immun. 1985;48(2):446–51. doi: 10.1128/iai.48.2.446-451.1985. PubMed PMID: 3988342; PubMed Central PMCID: PMCPMC261341.

61. Gundogdu O, Mills DC, Elmi A, Martin MJ, Wren BW, Dorrell N. The *Campylobacter jejuni* Transcriptional Regulator Cj1556 Plays a Role in the Oxidative and Aerobic Stress Response and Is Important for Bacterial Survival In Vivo. Journal of Bacteriology. 2011;193(16):4238–49. doi: doi:10.1128/jb.05189-11.

62. Gundogdu O, da Silva DT, Mohammad B, Elmi A, Mills DC, Wren BW, et al. The *Campylobacter jejuni* MarR-like transcriptional regulators RrpA and RrpB both influence bacterial responses to oxidative and aerobic stresses. Frontiers in Microbiology. 2015;Volume 6 -2015. doi: 10.3389/fmicb.2015.00724.

63. Srinivasan B, Kolli AR, Esch MB, Abaci HE, Shuler ML, Hickman JJ. TEER measurement techniques for in vitro barrier model systems. J Lab Autom. 2015;20(2):107–26. Epub 20150113. doi: 10.1177/2211068214561025. PubMed PMID: 25586998; PubMed Central PMCID: PMCPMC4652793.

64. Gerkins C, Hajjar R, Oliero M, Santos MM. Assessment of Gut Barrier Integrity in Mice Using Fluorescein-Isothiocyanate-Labeled Dextran. J Vis Exp. 2022;(189). Epub 20221118. doi: 10.3791/64710. PubMed PMID: 36468715.

65. MacCallum A, Hardy SP, Everest PH. *Campylobacter jejuni* inhibits the absorptive transport functions of Caco-2 cells and disrupts cellular tight junctions. Microbiology (Reading). 2005;151(Pt 7):2451–8. doi: 10.1099/mic.0.27950-0. PubMed PMID: 16000735.

66. Hofreuter D, Tsai J, Watson RO, Novik V, Altman B, Benitez M, et al. Unique features of a highly pathogenic Campylobacter jejuni strain. Infect Immun. 2006;74(8):4694–707. doi: 10.1128/IAI.00210-06. PubMed PMID: 16861657; PubMed Central PMCID: PMCPMC1539605.

67. Mills DC, Gundogdu O, Elmi A, Bajaj-Elliott M, Taylor PW, Wren BW, et al. Increase in *Campylobacter jejuni* invasion of intestinal epithelial cells under low-oxygen coculture conditions that reflect the in vivo environment. Infect Immun. 2012;80(5):1690–8. Epub 20120221. doi: 10.1128/iai.06176-11. PubMed PMID: 22354027; PubMed Central PMCID: PMCPMC3347453.

68. Brás AM, Ketley JM. Transcellular translocation of *Campylobacter jejuni* across human polarised epithelial monolayers. FEMS Microbiology Letters. 1999;179(2):209–15. doi: 10.1111/j.1574-6968.1999.tb08729.x.

69. Konkel ME, Mead DJ, Hayes SF, Cieplak W, Jr. Translocation of *Campylobacter jejuni* across human polarized epithelial cell monolayer cultures. J Infect Dis. 1992;166(2):308–15. doi: 10.1093/infdis/166.2.308. PubMed PMID: 1634802.

70. Kalischuk LD, Inglis GD, Buret AG. *Campylobacter jejuni* induces transcellular translocation of commensal bacteria via lipid rafts. Gut Pathog. 2009;1(1):2. Epub 20090203. doi: 10.1186/1757-4749-1-2. PubMed PMID: 19338680; PubMed Central PMCID: PMCPMC2653720.

71. Kalischuk LD, Leggett F, Inglis GD. *Campylobacter jejuni* induces transcytosis of commensal bacteria across the intestinal epithelium through M-like cells. Gut Pathogens. 2010;2(1):14. doi: 10.1186/1757-4749-2-14.

72. Lara-Tejero M, Galán JE. A bacterial toxin that controls cell cycle progression as a deoxyribonuclease I-like protein. Science. 2000;290(5490):354–7. doi: 10.1126/science.290.5490.354. PubMed PMID: 11030657.

73. De Rycke J, Oswald E. Cytolethal distending toxin (CDT): a bacterial weapon to control host cell proliferation? FEMS Microbiol Lett. 2001;203(2):141–8. doi: 10.1111/j.1574-6968.2001.tb10832.x. PubMed PMID: 11583839.

74. Gu J, Lin Y, Wang Z, Pan Q, Cai G, He Q, et al. *Campylobacter jejuni* Cytolethal Distending Toxin Induces GSDME-Dependent Pyroptosis in Colonic Epithelial Cells. Frontiers in Cellular and Infection Microbiology. 2022;12. doi: 10.3389/fcimb.2022.853204.

75. Fox James G, Rogers Arlin B, Whary Mark T, Ge Z, Taylor Nancy S, Xu S, et al. Gastroenteritis in NF-κB-Deficient Mice Is Produced with Wild-Type *Camplyobacter jejuni* but Not with *C. jejuni* Lacking Cytolethal Distending Toxin despite Persistent Colonization with Both Strains. Infection and Immunity. 2004;72(2):1116–25. doi: 10.1128/iai.72.2.1116-1125.2004.

76. Jain D, Prasad KN, Sinha S, Husain N. Differences in virulence attributes between cytolethal distending toxin positive and negative *Campylobacter jejuni* strains. Journal of Medical Microbiology. 2008;57(3):267–72. doi: 10.1099/jmm.0.47317-0.

77. Pickett CL, Pesci EC, Cottle DL, Russell G, Erdem AN, Zeytin H. Prevalence of cytolethal distending toxin production in *Campylobacter jejuni* and relatedness of *Campylobacter* sp. cdtB gene. Infect Immun. 1996;64(6):2070–8. doi: 10.1128/iai.64.6.2070-2078.1996. PubMed PMID: 8675309; PubMed Central PMCID: PMCPMC174038.

78. Talukder KA, Aslam M, Islam Z, Azmi IJ, Dutta DK, Hossain S, et al. Prevalence of virulence genes and cytolethal distending toxin production in *Campylobacter jejuni* isolates from diarrheal patients in Bangladesh. J Clin Microbiol. 2008;46(4):1485–8. Epub 20080220. doi: 10.1128/jcm.01912-07. PubMed PMID: 18287317; PubMed Central PMCID: PMCPMC2292937.

79. Eyigor A, Dawson KA, Langlois BE, Pickett CL. Cytolethal distending toxin genes in *Campylobacter jejuni* and *Campylobacter coli* isolates: detection and analysis by PCR. J Clin Microbiol. 1999;37(5):1646–50. doi: 10.1128/jcm.37.5.1646-1650.1999. PubMed PMID: 10203548; PubMed Central PMCID: PMCPMC84865.

80. Hickey TE, Baqar S, Bourgeois AL, Ewing CP, Guerry P. *Campylobacter jejuni-*stimulated secretion of interleukin-8 by INT407 cells. Infect Immun. 1999;67(1):88–93. doi: 10.1128/IAI.67.1.88-93.1999. PubMed PMID: 9864200; PubMed Central PMCID: PMCPMC96281.

81. Zheng J, Meng J, Zhao S, Singh R, Song W. *Campylobacter*-induced interleukin-8 secretion in polarized human intestinal epithelial cells requires *Campylobacter*-secreted cytolethal distending toxin- and Toll-like receptor-mediated activation of NF-kappaB. Infect Immun. 2008;76(10):4498–508. Epub 20080721. doi: 10.1128/IAI.01317-07. PubMed PMID: 18644884; PubMed Central PMCID: PMCPMC2546826.

82. Hickey TE, McVeigh AL, Scott DA, Michielutti RE, Bixby A, Carroll SA, et al. *Campylobacter jejuni* cytolethal distending toxin mediates release of interleukin-8 from intestinal epithelial cells. Infect Immun. 2000;68(12):6535–41. doi: 10.1128/IAI.68.12.6535-6541.2000. PubMed PMID: 11083762; PubMed Central PMCID: PMCPMC97747.

83. MacCallum AJ, Harris D, Haddock G, Everest PH. *Campylobacter jejuni*-infected human epithelial cell lines vary in their ability to secrete interleukin-8 compared to in vitro-infected primary human intestinal tissue. Microbiology (Reading). 2006;152(Pt 12):3661–5. doi: 10.1099/mic.0.29234-0. PubMed PMID: 17159219.

84. Van Deun K, Haesebrouck F, Heyndrickx M, Favoreel H, Dewulf J, Ceelen L, et al. Virulence properties of *Campylobacter jejuni* isolates of poultry and human origin. J Med Microbiol. 2007;56(Pt 10):1284–9. doi: 10.1099/jmm.0.47342-0. PubMed PMID: 17893162.

85. Malik-Kale P, Raphael BH, Parker CT, Joens LA, Klena JD, Quiñones B, et al. Characterization of genetically matched isolates of *Campylobacter jejuni* reveals that mutations in genes involved in flagellar biosynthesis alter the organism’s virulence potential. Appl Environ Microbiol. 2007;73(10):3123–36. Epub 20070316. doi: 10.1128/aem.01399-06. PubMed PMID: 17369342; PubMed Central PMCID: PMCPMC1907099.

86. Korlath JA, Osterholm MT, Judy LA, Forfang JC, Robinson RA. A point-source outbreak of campylobacteriosis associated with consumption of raw milk. J Infect Dis. 1985;152(3):592–6. doi: 10.1093/infdis/152.3.592. PubMed PMID: 4031557.

87. Bacon DJ, Alm RA, Burr DH, Hu L, Kopecko DJ, Ewing CP, et al. Involvement of a plasmid in virulence of *Campylobacter jejuni* 81-176. Infect Immun. 2000;68(8):4384–90. doi: 10.1128/IAI.68.8.4384-4390.2000. PubMed PMID: 10899834; PubMed Central PMCID: PMCPMC98329.

88. Pei Z, Burucoa C, Grignon B, Baqar S, Huang XZ, Kopecko DJ, et al. Mutation in the peb1A locus of *Campylobacter jejuni* reduces interactions with epithelial cells and intestinal colonization of mice. Infect Immun. 1998;66(3):938–43. doi: 10.1128/iai.66.3.938-943.1998. PubMed PMID: 9488379; PubMed Central PMCID: PMCPMC107999.

89. Black RE, Levine MM, Clements ML, Hughes TP, Blaser MJ. Experimental *Campylobacter jejuni* infection in humans. J Infect Dis. 1988;157(3):472–9. doi: 10.1093/infdis/157.3.472. PubMed PMID: 3343522.

90. Corcionivoschi N, Clyne M, Lyons A, Elmi A, Gundogdu O, Wren BW, et al. *Campylobacter jejuni* Cocultured with Epithelial Cells Reduces Surface Capsular Polysaccharide Expression. Infection and Immunity. 2009;77(5):1959–67. doi: doi:10.1128/iai.01239-08.

91. Corcionivoschi N, Clyne M, Lyons A, Elmi A, Gundogdu O, Wren BW, et al. *Campylobacter jejuni* cocultured with epithelial cells reduces surface capsular polysaccharide expression. Infect Immun. 2009;77(5):1959–67. Epub 20090309. doi: 10.1128/iai.01239-08. PubMed PMID: 19273563; PubMed Central PMCID: PMCPMC2681765.

92. Taboada EN, Acedillo RR, Carrillo CD, Findlay WA, Medeiros DT, Mykytczuk OL, et al. Large-scale comparative genomics meta-analysis of *Campylobacter jejuni* isolates reveals low level of genome plasticity. J Clin Microbiol. 2004;42(10):4566–76. doi: 10.1128/jcm.42.10.4566-4576.2004. PubMed PMID: 15472310; PubMed Central PMCID: PMCPMC522315.

93. Woodcock DJ, Krusche P, Strachan NJC, Forbes KJ, Cohan FM, Méric G, et al. Genomic plasticity and rapid host switching can promote the evolution of generalism: a case study in the zoonotic pathogen *Campylobacter*. Scientific Reports. 2017;7(1):9650. doi: 10.1038/s41598-017-09483-9.

94. Hofreuter D, Tsai J, Watson Robert O, Novik V, Altman B, Benitez M, et al. Unique Features of a Highly Pathogenic *Campylobacter jejuni* Strain. Infection and Immunity. 2006;74(8):4694–707. doi: 10.1128/iai.00210-06.

95. Leonard EE, 2nd, Tompkins LS, Falkow S, Nachamkin I. Comparison of *Campylobacter jejuni* isolates implicated in Guillain-Barré syndrome and strains that cause enteritis by a DNA microarray. Infect Immun. 2004;72(2):1199–203. doi: 10.1128/iai.72.2.1199-1203.2004. PubMed PMID: 14742576; PubMed Central PMCID: PMCPMC321608.

96. Parker Craig T, Quiñones B, Miller William G, Horn Sharon T, Mandrell Robert E. Comparative Genomic Analysis of *Campylobacter jejuni* Strains Reveals Diversity Due to Genomic Elements Similar to Those Present in C. jejuni Strain RM1221. Journal of Clinical Microbiology. 2006;44(11):4125–35. doi: 10.1128/jcm.01231-06.

