## Supplementary Figures for "Strain-specific virulence signatures of *Campylobacter jejuni* associated with watery versus bloody diarrhea in the neonatal piglet model"

Supplementary Figure S1

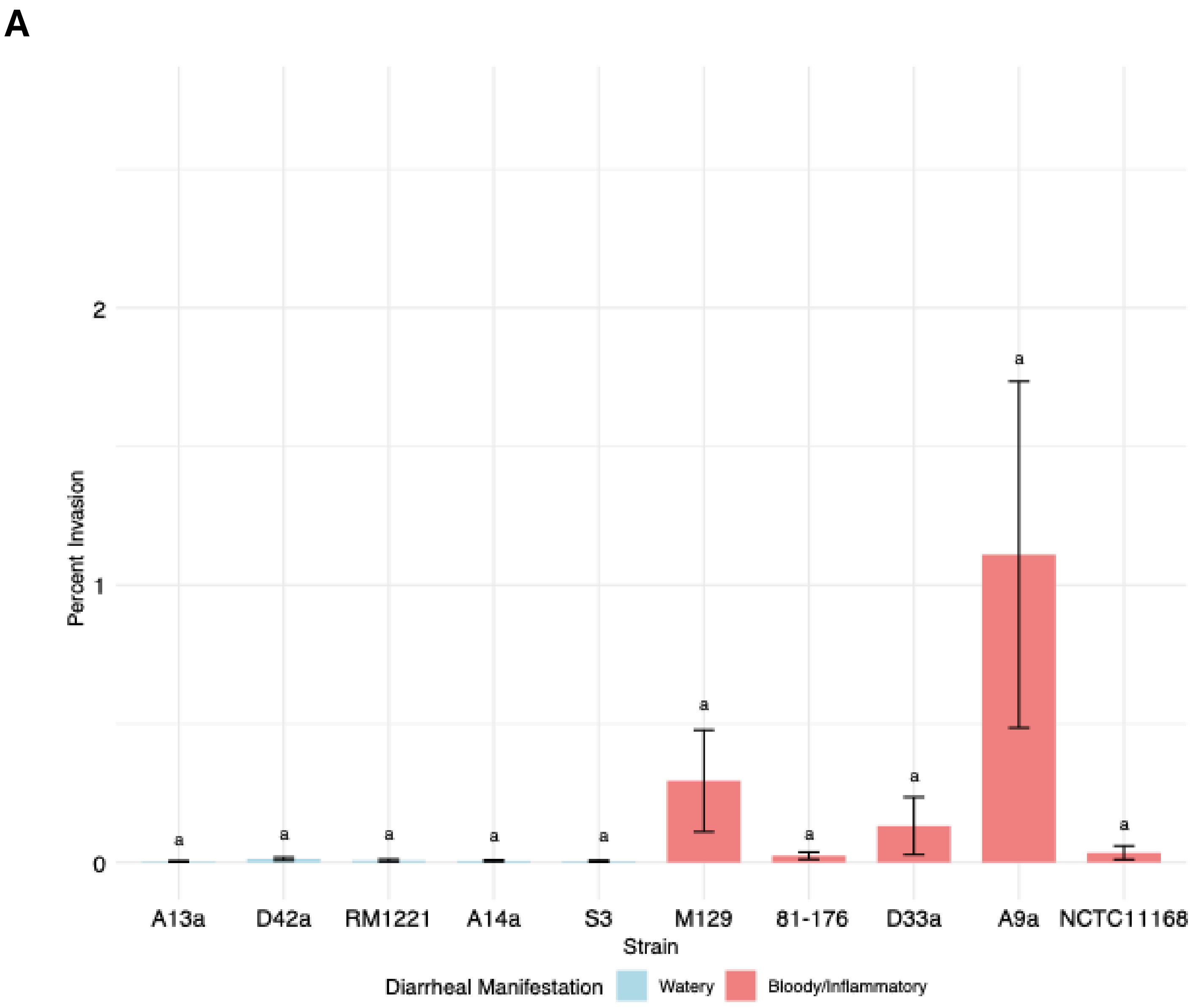


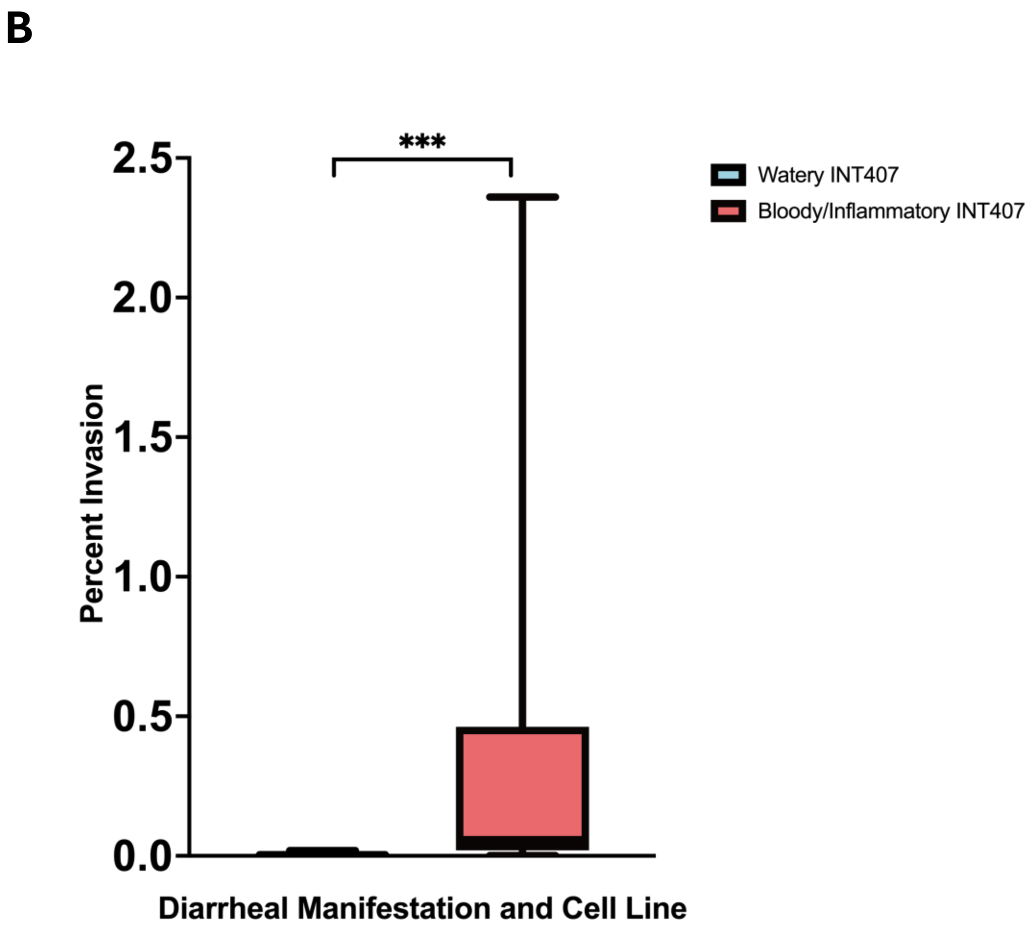

Supplementary Figure S2

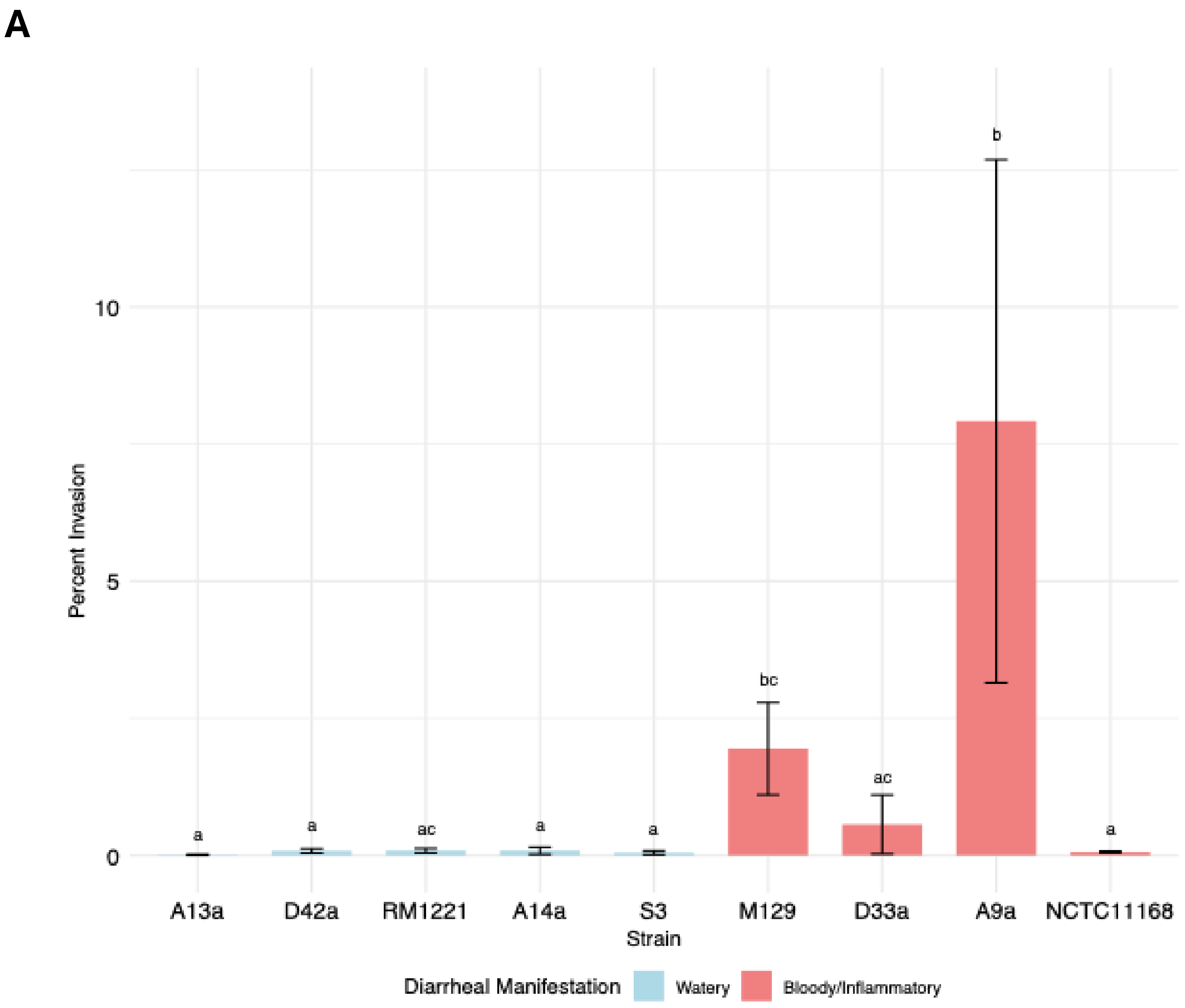


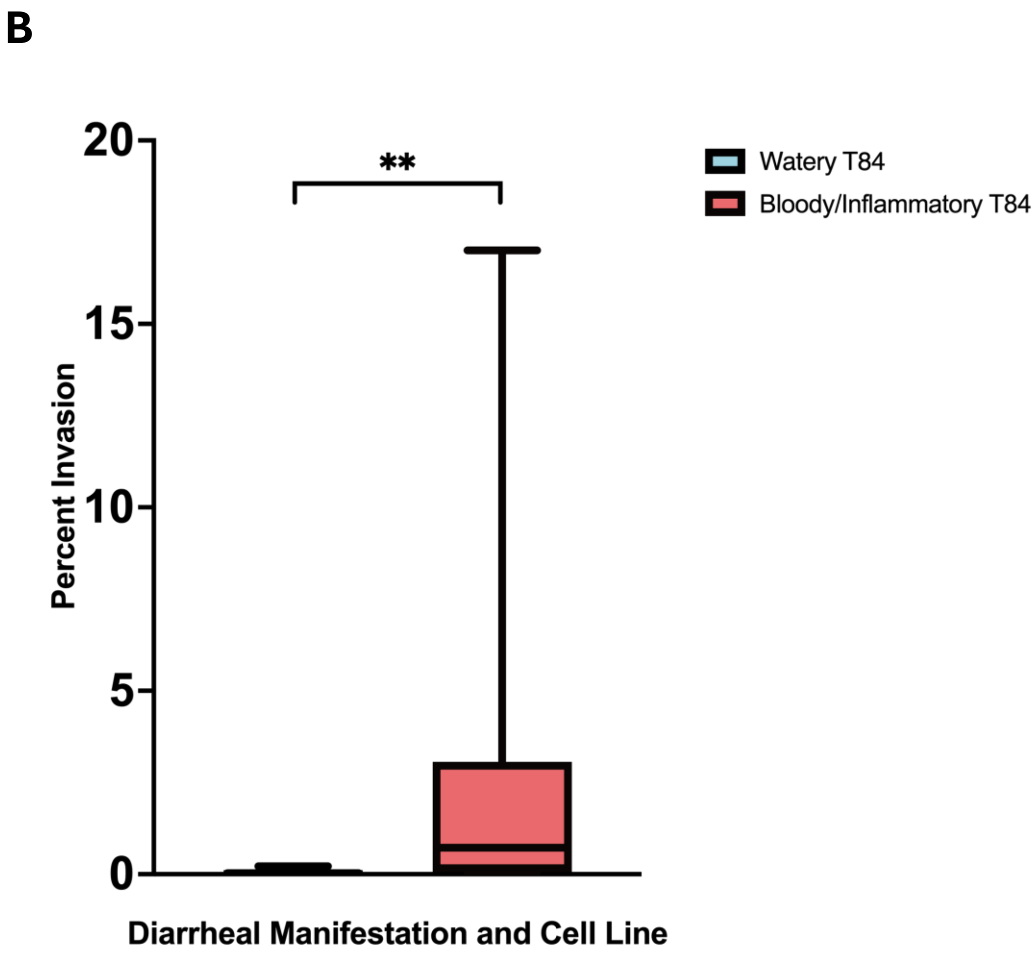

Supplementary Figure S3

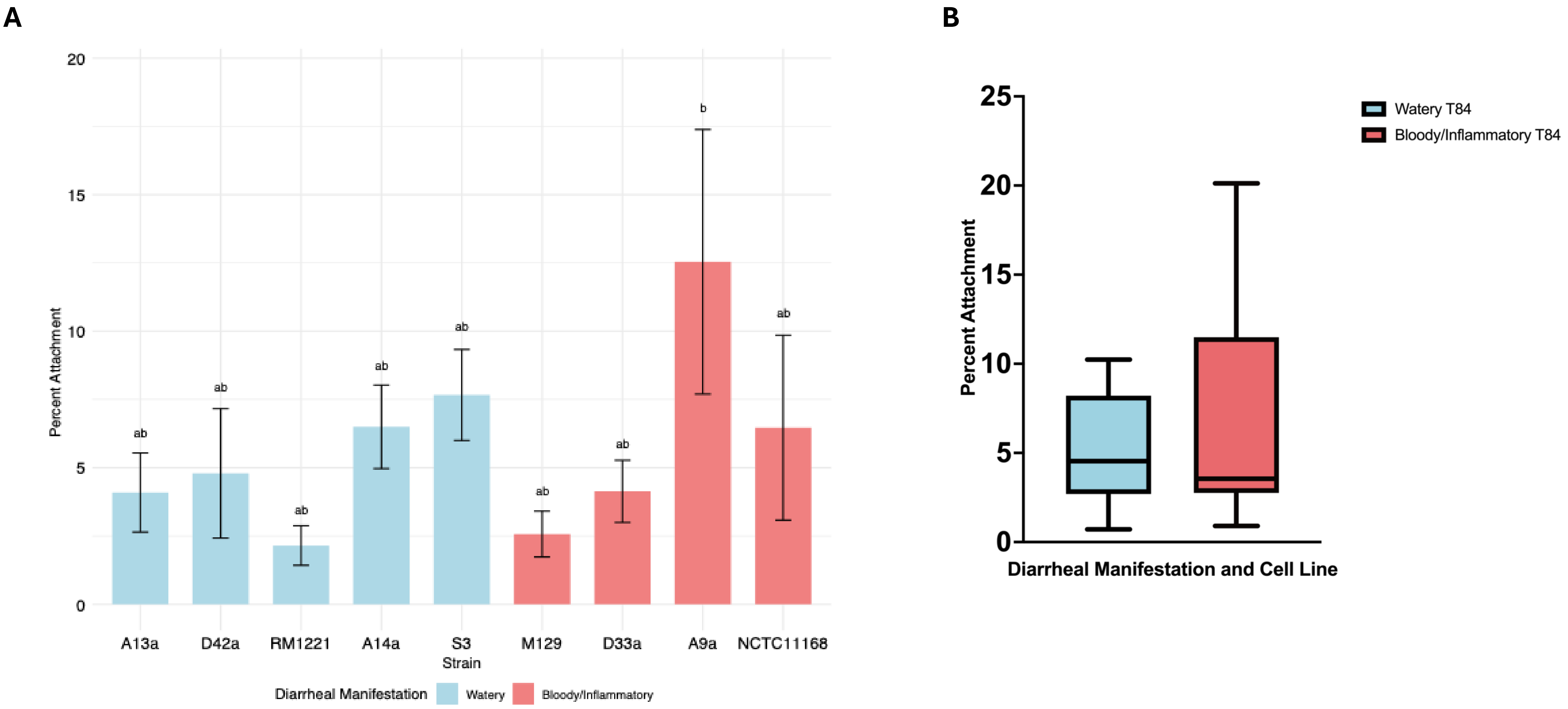

Supplementary Figure S4

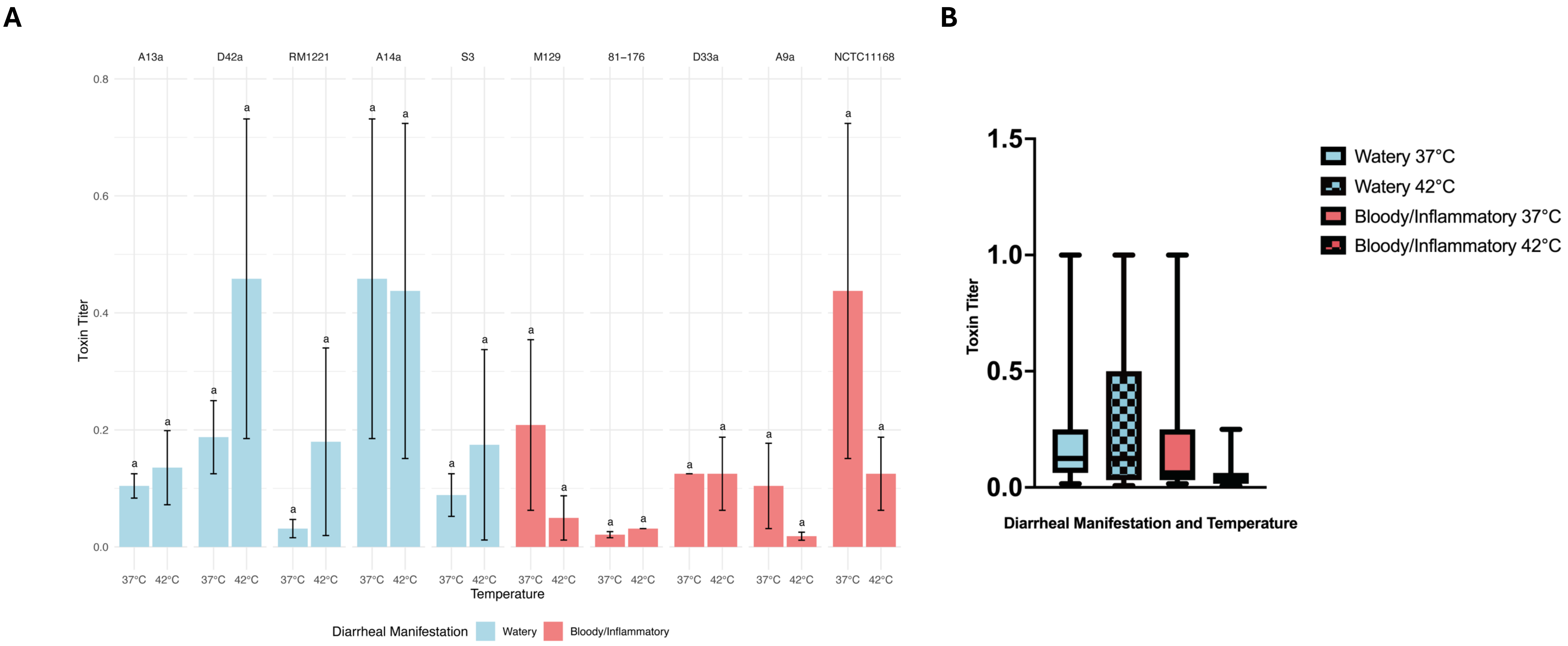

Supplementary Figure S5

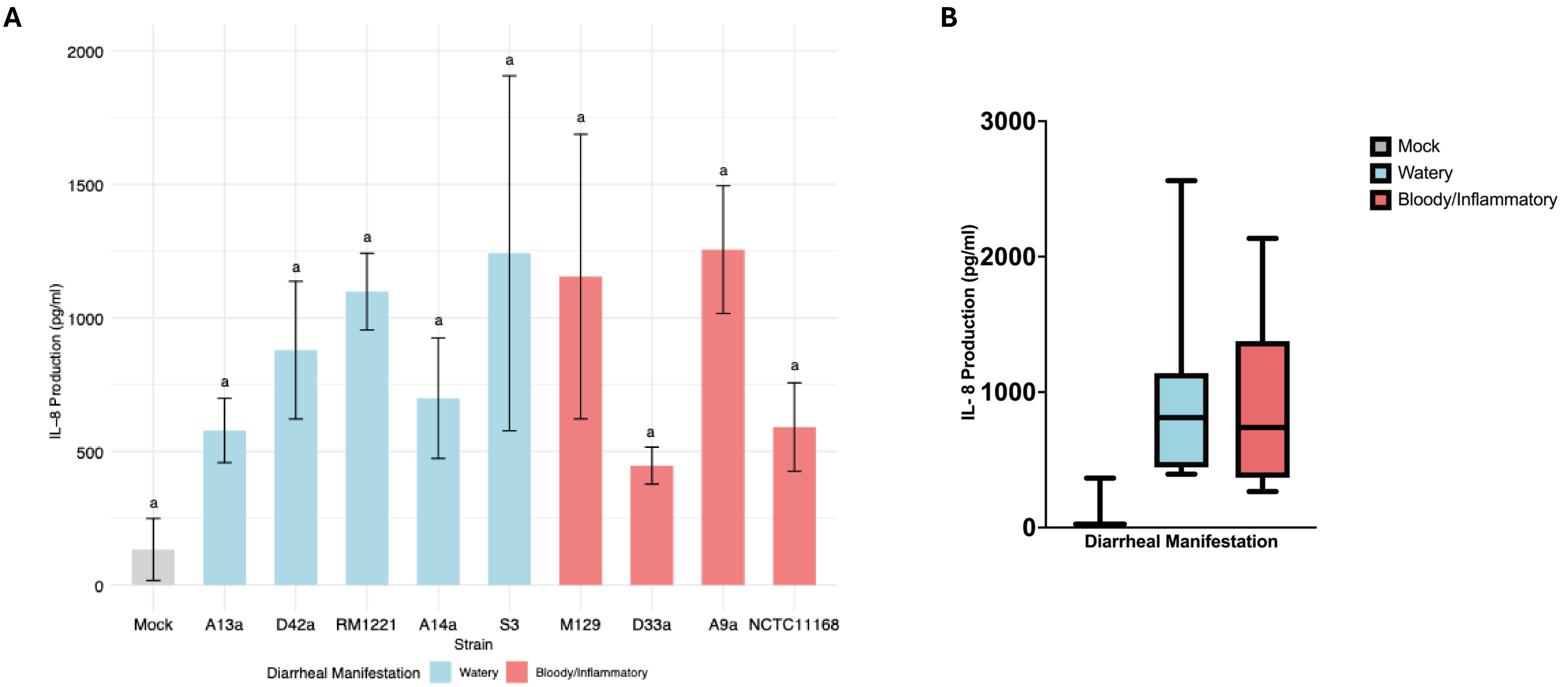

Supplementary Figure S6

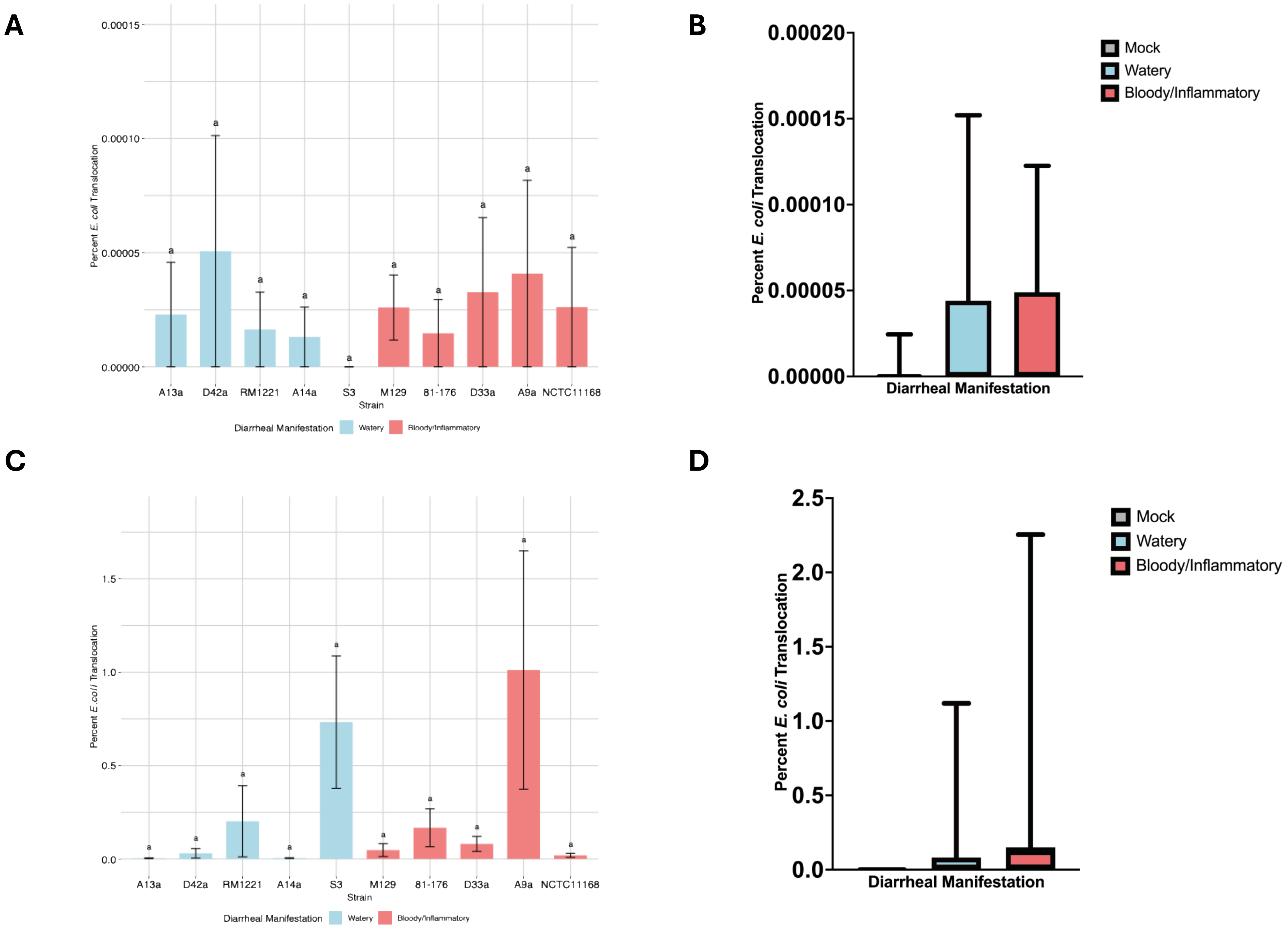
